# Generalization but not strengthening of negative memories drives the development of depression-like behaviors

**DOI:** 10.1101/2025.02.10.637591

**Authors:** Xin Cheng, Yubo Hu, Yan Zhao, Panwu Zhao, Xiaomeng Bai, Yi Chen, Deshan Kong, Shuyu Zheng, Yuena Zheng, Yumeng Wang, Yanni Zeng, Wei-Jye Lin, Xiaojing Ye

## Abstract

Individuals with depression exhibit intensified and generalized negative memories. However, how prior memories actually influence onset and/or progression of depression, particularly whether the strengthening or the generalization of prior memories play a key role, is poorly understood. Here, we introduced behavioral paradigms to differentiate between memory strengthening and generalization, and found that negative experiences that produced memory overgeneralization, but not strengthening, induced depression-like behaviors. Furthermore, we identified that the medial prefrontal cortex (mPFC) to the bed nucleus of the stria terminals (BNST) projection functionally linked memory generalization to depression, together with single-cell calcium imaging which revealed a significant overlap in the mPFC^BNST^ neuronal ensemble encoding depression-like behaviors with that encoding generalized memory, but not memory strength. Circuit-based transcriptomic analysis and chromophore-assisted light inactivation demonstrated that triggering actin remodeling in the mPFC^BNST^ neurons, a memory consolidation mechanism enhanced generalization while reducing memory strength, also induced depression-like behaviors. Collectively, these findings suggest generalization of negative memories as a primary factor that drives depression-like behaviors, highlighting the importance of early identification of individuals at risk for depression who exhibit overgeneralized negative memories, and targeting overgeneralized negative memories in the treatment of depression.

## INTRODUCTION

Depression is a prevalent and debilitating mental disorder^1^. Despite the availability of various antidepressants that can alleviate depressive symptoms to some extent, their efficacy in altering the progression of the disorder is limited and the recurrence rate remains high^2^. Therefore, there is an urgent need to address the underlying mechanisms and improve long-term outcomes with the development of effective strategies that can genuinely slow or reverse the trajectory of depression.

A specific cognitive model of depression hypothesized that negative experiences create negative cognitive biases within individuals, leading to depression^3^. Consequently, therapeutic interventions aimed at modifying these cognitive biases hold considerable clinical potential^3^. This theoretical framework is bolstered by extensive clinical reports documenting distortions of autobiographical memories among depressed individuals. Specifically, individuals with depression tend to frequently recall negative memories, which are not only vivid but emotionally charged, thereby triggering repetitive negative thoughts associated with greater risk of depression^4, 5^. Furthermore, they often generalize negative experiences associated with specific contexts to a broader sense of self-worth, which excessively controls over their behaviors even in neutral environments and subsequently provoking depressive symptoms^6–8^. However, despite these observations, the specific components of negative memories that influence the development of depression and the underlying neurobiological mechanisms remain poorly understood. Indeed, within the realm of depression research, most studies have viewed memory alterations in depressed individuals as cognitive symptoms arising from stress, rather than as contributory factors to the disorder itself ^9–11^.

Memory, a fundamental cognitive function of the brain, has two key components: strength, which refers to the intensity of recalled memories, and generalization, the process of applying memories to comparable yet distinct situations. The field of memory research has predominantly focused on memory strength and its clinical relevance^12–16^. In contrast, generalization has often been undervalued, primarily regarded as a parametric feature of memories. The impact of negative memory generalization on neuropsychiatric disorders, particularly depression, warrants further investigation. It remains unclear whether the strengthening or generalization of negative memory plays a pivotal role in the onset and progression of depression. Identifying the key component of negative memories that links to depression is essential for the early detection of at-risk individuals and for determining whether reducing the strength or limiting the generalization of negative memories should be the primary focus of therapies aimed at mitigating the effects of negative cognitive bias on depression^13–15, 17^. Such insights are critical for developing effective treatments for depression.

Memory strengthening and generalization are often intertwined, with stronger negative memories typically accompanied by increased generalization^18, 19^. However, there is a scarcity of experimental models that can clearly distinguish between these two key components of memory. In this study, we introduced behavioral paradigms that clearly dissociated memory strengthening and generalization. Unexpectedly, our results indicate that the generalization of negative memory, rather than its strengthening as traditionally thought, is a principal driver for depression-like behaviors. These findings highlight the importance of rectifying overgeneralization of negative memories and its related mechanisms as potential therapeutic approaches for depression and other stress-related neuropsychiatric disorders.

## MATERIALS AND METHODS

The detailed methods for the mice, behavioral training and testing, stereotaxic surgery, circuit manipulations, immunofluorescence staining and image analysis, single-cell calcium imaging, Retro-TRAP and RNA sequencing, statistical analyses can be found in Supplementary Materials and Methods.

## RESULTS

### Generalization, but not strengthening, of negative memories are associated with depression-like behaviors

To explore whether the strength or generalization of negative memories is connected to depression, we needed to establish new behavioral paradigms that could distinguish these two aspects of memory. Towards that end, we examined how the number and variability of negative experiences affected the strength and generalization of memories. We trained mice with single contextual fear conditioning (CFC) training (referred to as “1xTr”), or two CFC training sessions in either the same (“2xTr same”) or different contexts (“2xTr altered”). Memory strength was measured in the same context as training, and memory generalization was measured in a modified context largely deviated from training (**Fig. 1A**).

**Fig. 1.**
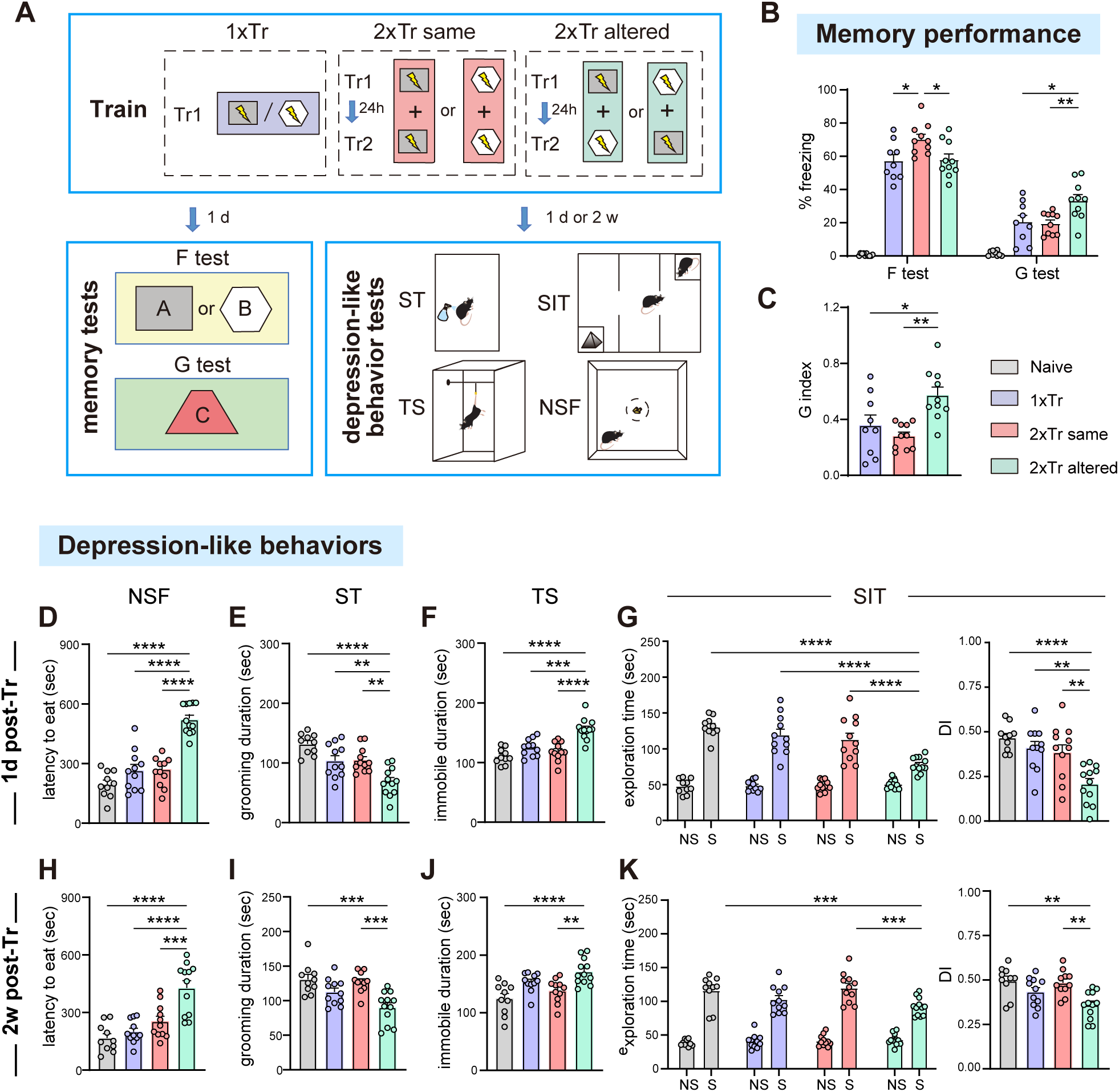
Depression-like behaviors are associated with the generalization, but not the strengthening, of negative memories. (**A**) Schematics of contextual fear conditioning (CFC) training (Tr), fear memory strength test (F test) and fear generalization test (G test), novelty-suppressed feeding (NSF) test, splash test (ST), tail suspension (TS) test and social interaction test (SIT). Naïve: control mice without training; 1xTr: a single CFC training; 2xTr same: two CFC training trials in the same context; 2xTr altered: two CFC training trials in two different contexts. (**B-C**), Quantification of the percentage of time spent in freezing during F test and G test (B), and the generalization index (G index, C). G index: % freezing during G test divided by % freezing during F test. n = 9-10 per group. (**D, H**) Quantification of the latency to eat in NSF test. (**E, I**) Quantification of grooming duration in ST. (**F, J**) Quantification of the immobile time during TS test. (**G, K**) Quantification of time exploring the non-social target (NS) and the social mouse (S) during SIT, as well as the differential index (DI) during SIT. DI: time exploring the stranger mouse subtracted by time exploring the toy mouse as a fraction of total exploration time. D-G, tests starting 1 d after training. H-K, tests starting 2 w after training. D-K, n = 10-12 per group. Data are presented as mean ± s.e.m. and analyzed by one-way ANOVA followed by Turkey’s post hoc test or two-way ANOVA followed the Sidak’s *post hoc* test. * *p* < 0.05, ** *p* < 0.01, *** *p* < 0.001, **** *p* < 0.0001.

Compared to “1xTr”, the “2xTr same” paradigm significantly enhanced memory strength in mice of both sexes, whereas the “2xTr altered” paradigm resulted in memory strength similar to “1xTr” (**Fig. 1B**). During memory generalization test, mice in the “1xTr” and “2xTr same” groups exhibited similarly low levels of freezing, whereas the “2xTr altered” paradigm significantly enhanced freezing level in the memory generalization test (**Fig. 1B**). Since memory strength has an impact on the outcome of memory generalization test, we also calculated the generalization index (G index), which was derived from the ratio of freezing in the generalization test to that in the memory strength test to offer a more robust measure of memory generalization^19, 20^. Compared to the “1xTr” and “2xTr same” groups, the “2xTr altered” group also exhibited a significantly increased G index (**Fig. 1C**). These results suggest that a second training in the same versus different contexts induced memory strengthening and generalization, respectively, providing a double dissociation between the two components of memory.

We next evaluated the ability of different patterns of CFC training to induce depression-like behaviors. We found that during the depression-like behavioral tests starting one day after training, mice in the “2xTr altered”, but not the “1xTr” or “2xTr same” group, exhibited significant depression-like phenotypes across various tests. These included increased latency to eat in the novelty-suppressed feeding test as an indication of reduced motivation, decreased grooming duration in the splash test as an indication of apathy, increased immobility in the tail suspension test as an indication of helplessness, and decreased social preference in the social interaction test as an indication of social withdrawal and anhedonia (**Fig. 1D-G**). Notably, these behavioral phenotypes persisted for at least two weeks after training, aligning with clinical diagnostic criteria for depression based on the duration of symptoms (**Fig. 1H-K**). Additionally, in a subset of animals that underwent both memory and depression-like behavior assessments, we examined the correlation between memory performances and depression-like behaviors. Our analysis revealed significant correlations of depression-like behaviors with indicators of memory generalization, including the freezing level during generalization test and the G index. In contrast, no significant correlation was observed between depression-like behaviors and freezing level during the memory strength test (**Supplementary Fig. 1**).

Taken together, by differentiating between memory strengthening and generalization, our results reveal unexpectedly that negative memory generalization, rather than its strengthening, is behaviorally associated with the occurrence of depression-like behaviors.

### Activation of the mPFC-to-BNST projection promotes the generalization, but not strengthening of negative memories

The unpredictability of negative experiences is recognized as a crucial factor in the onset of depression^21^. Yet, the distinctions between memories of unpredictable versus predictable negative events, and their impact on depressive behaviors, remain elusive. Given that the “2xTr altered” versus “2xTr same” paradigms induced memory generalization and strengthening, respectively, we postulate that memory generalization might be a critical mechanism linking unpredictable negative experiences to depression.

To identify the neural circuit underpinning memory generalization but not strengthening, we performed a brain-wide mapping of c-FOS after subjecting mice to different patterns of CFC training, to identify features in brain networks recruited by training patterns that induced negative memory generalization versus strengthening (**Fig. 2A**). Since multiple brain regions were activated by CFC trainings, we computed the interregional correlations of the c-FOS^+^ cell density to assess functional connectivity, from which we generated networks of co-regulated brain regions (**Fig. 2B-D, Supplementary Table 1**)^22, 23^. Compared to other conditions, the “2xTr altered” condition exhibited a much denser connectivity, indicating increased interregional crosstalk (**Fig. 2B-D**). By calculating changes in the network global efficiency after isolating each node^23, 24^, we identified the bed nucleus of the stria terminalis (BNST) as the top hub for network recruited in “2xTr altered” paradigm (**Fig. 2D**), which led to generalized CFC memory.

**Fig. 2.**
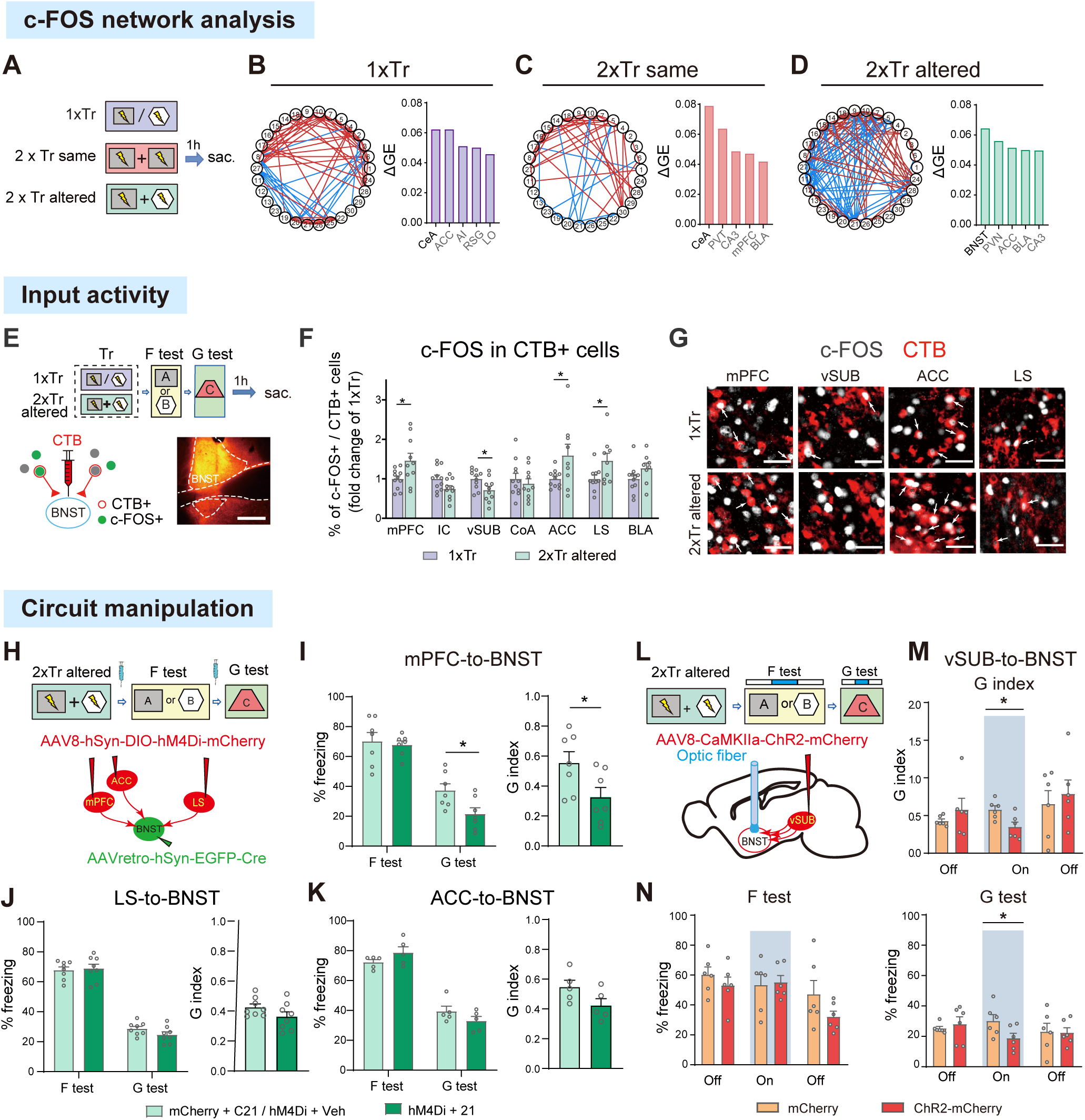
The mPFC-to-BNST projection are activated and required for negative memory generalization following variable negative experiences. (**A**) Schematic of CFC training and brain collection (sac). (**B-D**) Left: network graphs derived from c-FOS mapping, with nodes representing each brain region (see Table S1 for the list) and the edges representing significant correlations (Pearson’s *p* < 0.05; blue: negative correlation; red: positive correlation). Right: top 5 brain regions that contribute to network global efficiency. n = 7-9 per group. (**E**) Schematic of the experiment timeline, as well as cholera toxin B (CTB) injection into BNST and co-staining with c-FOS. A representative image of CTB infusion into BNST. Scale bar, 350 µm. (**F**) Fold changes in c-FOS expression in the CTB^+^ BNST-projecting neurons in upstream regions induced by the G test, normalized to the mean value for the “1xTr” group in each region. mPFC: medial prefrontal cortex; IC: insular cortex; vSUB: ventral subiculum; CoA: anterior cortical amygdala; ACC: anterior cingulate cortex; LS: lateral septum; BLA: basolateral amygdala. (**G**) Representative images of c-FOS staining (grey) in the CTB-labelled BNST-projecting neurons (red) in various upstream brain regions. Scale bar, 40 µm. n = 8-11 per group. (**H, L**) Schematics of the behavioral paradigm and AAV infusion for circuit-specific manipulation. (**I-K**) Left: quantification of the percentage of time spent in freezing during fear memory strength test (F test) and generalization (G test) for chemogenetic inhibition of different projections into the BNST during memory tests after the “2xTr altered” paradigm. Right: the generalization index (G index), calculated as % freezing during G test divided by % freezing during F test. n = 7 (I, mPFC-to-BNST), 8 (J, LS-to-BNST), 5 (K, ACC-to-BNST) per group. (**M**) Quantification of the G index during light on/off phases for optogenetic activation of the vSUB-to-BNST projection. (**N**) Quantification of the percentage of time spent in freezing during F and G tests for optogenetic activation of the vSUB-to-BNST projection. n = 6 per group. Data are presented as mean ± s.e.m. and analyzed by Student’s *t* tests or two-way ANOVA followed by Sidak’s *post hoc* test. * *p* < 0.05.

In contrast, the central amygdala (CeA) was identified as the top hub in the networks recruited in both the “1xTr” and “2xTr same” paradigms (**Fig. 2B-C**).

While the amygdala has been the focus in fear research, the BNST is a critical relay station connecting limbic and forebrain structures to the hypothalamic-pituitary-adrenal stress axis^25^. We then wondered whether the upstream inputs integrated by BNST prior to regulating negative memory expression were different in the memory strength test versus generalization test after “2xTr altered” paradigm. Combining retrograde tracing with c-FOS staining (**Fig. 2E**), our findings revealed that the enhanced memory generalization after the “2xTr altered” paradigm was associated with more activated BNST-projecting neurons in the medial prefrontal cortex (mPFC^BNST^), anterior cingulate cortex (ACC^BNST^) and lateral septum (LS^BNST^) but fewer in the ventral subiculum (vSUB^BNST^) (**Fig. 2F-G**). These patterns of activation were largely specific to BNST-projecting neurons in these regions, and did not apply to the whole investigated regions except for LS (**Supplementary Fig. 2A-C**). Also, they were not observed following the memory strength test (**Supplementary Fig. 2D-G**).

To examine the functional contribution of different upstream inputs to the BNST in memory expression after “2xTr altered”, we suppressed the activity of BNST-projecting neurons in the mPFC, ACC or LS during memory tests (**Fig. 2H**). Suppression of the mPFC^BNST^, but not ACC^BNST^ or LS^BNST^ neuronal activity significantly reduced freezing during the memory generalization test but not the memory strength test (**Fig. 2I-K**). Furthermore, consistent with the observation that memory generalization was accompanied by reduced activity in the vSUB^BNST^ neurons, optogenetic activation of these neurons significantly reduced memory generalization (**Fig. 2L-N, Supplementary Fig. 3A**).

The mPFC is known to play a critical role in both memory and depression ^26–29^. We further examined whether the mPFC-to-BNST projection played a role in generalization of fear memory under other conditions in addition to the “2xTr altered” condition. Memory generalization is known to be graded^30, 31^. Mice trained with “1xTr” showed significantly more freezing in a context similar to the training context (context A’) compared to a context that deviated greatly from training (context C) (**Fig. 3A**). Inhibition of the mPFC^BNST^ neurons significantly reduced generalization to the similar context (**Fig. 3B-C**). In the reverse fashion, following the “1xTr” or “2xTr same” paradigm in mice, optogenetic activation of the mPFC-to-BNST projection significantly enhanced the expression of fear memory generalization **(Fig. 3D-K, Supplementary Fig. 3B-C**). Of note, activation of the mPFC-to-BNST projection decreased freezing level during the memory strength test in the training context after “1xTr” **(Fig. 3E**), supporting the notion that memory generalization and strength can be oppositely regulated.

**Fig. 3.**
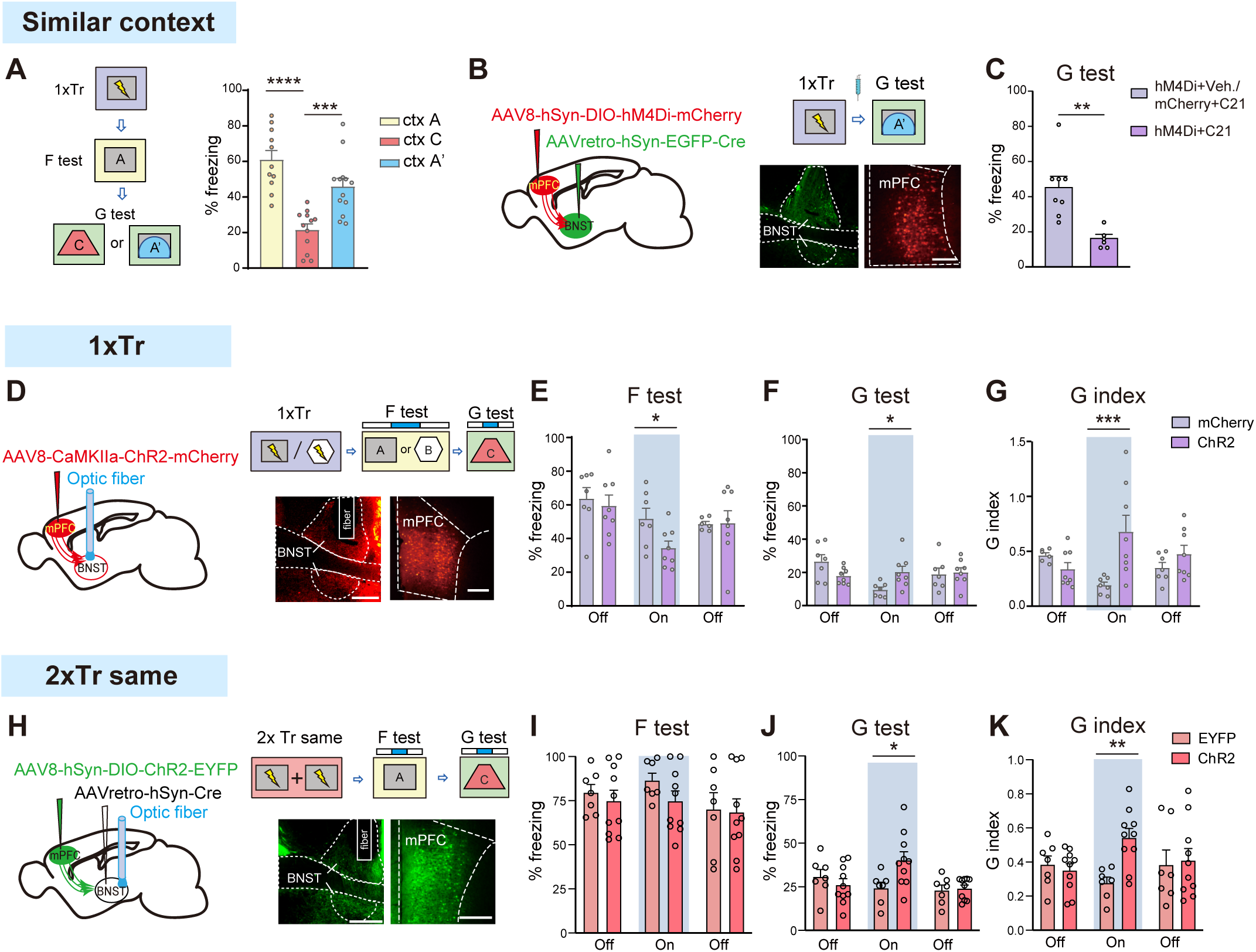
Activation of the mPFC-to-BNST projection promotes negative memory generalization, but not memory strengthening. (**A**) Left: Schematic of the experimental timeline. Right: Quantification of the percentage of time spent in freezing during memory tests in the same context as used in training (Ctx A), in a context largely deviated from the training context (Ctx C) or in a context similar to training (Ctx A’). n = 11-13 per group. (**B, D, H**) Schematics of AAV infusion, the experimental timeline and representative images of AAV-mediated fluorescent protein expression in the BNST and mPFC. Scale bar, 300 µm. (**C**) Quantification of changes in the percentage of time spent in freezing during the G test in context A’ by inhibition of the mPFC-to-BNST projection. n = 6-8 per group. (**E-F, I-J**) Quantification of the percentage of time spent in freezing in F test or G test, during light on/off phases for optogenetic activation of the mPFC-to-BNST projection after “1xTr” (E-F, n = 6-8 per group) or the “2xTr same” (I-J, n = 7-10 per group) paradigm. (**G, K**) Quantification of G index for optogenetic activation of the mPFC-to-BNST projection. Data are presented as mean ± s.e.m. and analyzed by Student’s *t* tests or two-way ANOVA followed by Sidak’s *post hoc* test. * *p* < 0.05, ** *p* < 0.01, *** *p* < 0.001, **** *p* < 0.0001.

Altogether, these findings suggest activation of mPFC-to-BNST projection as a critical neural circuit mechanism promoting the expression of generalized negative memories following variable negative experiences.

### The neuronal ensemble encoding negative memory generalization, but not strength, significantly overlaps with that encoding depression-like behavior

If memory generalization underlies the link between unpredictable negative experiences and depression, we postulated that there might exist overlapping neuronal ensembles encoding memory generalization and depression-like behaviors following unpredictable training. We utilized a miniaturized microscope to visualize Ca^2+^ activity in individual mPFC^BNST^ neurons in free-moving mice during consecutive behavioral assays, including two CFC trainings in the same or altered contexts, a memory strength test, a memory generalization test and a tail suspension test to evaluate helplessness, a core symptom of depression (**Fig. 4A-C, Supplementary Fig. 3D**).

**Fig. 4.**
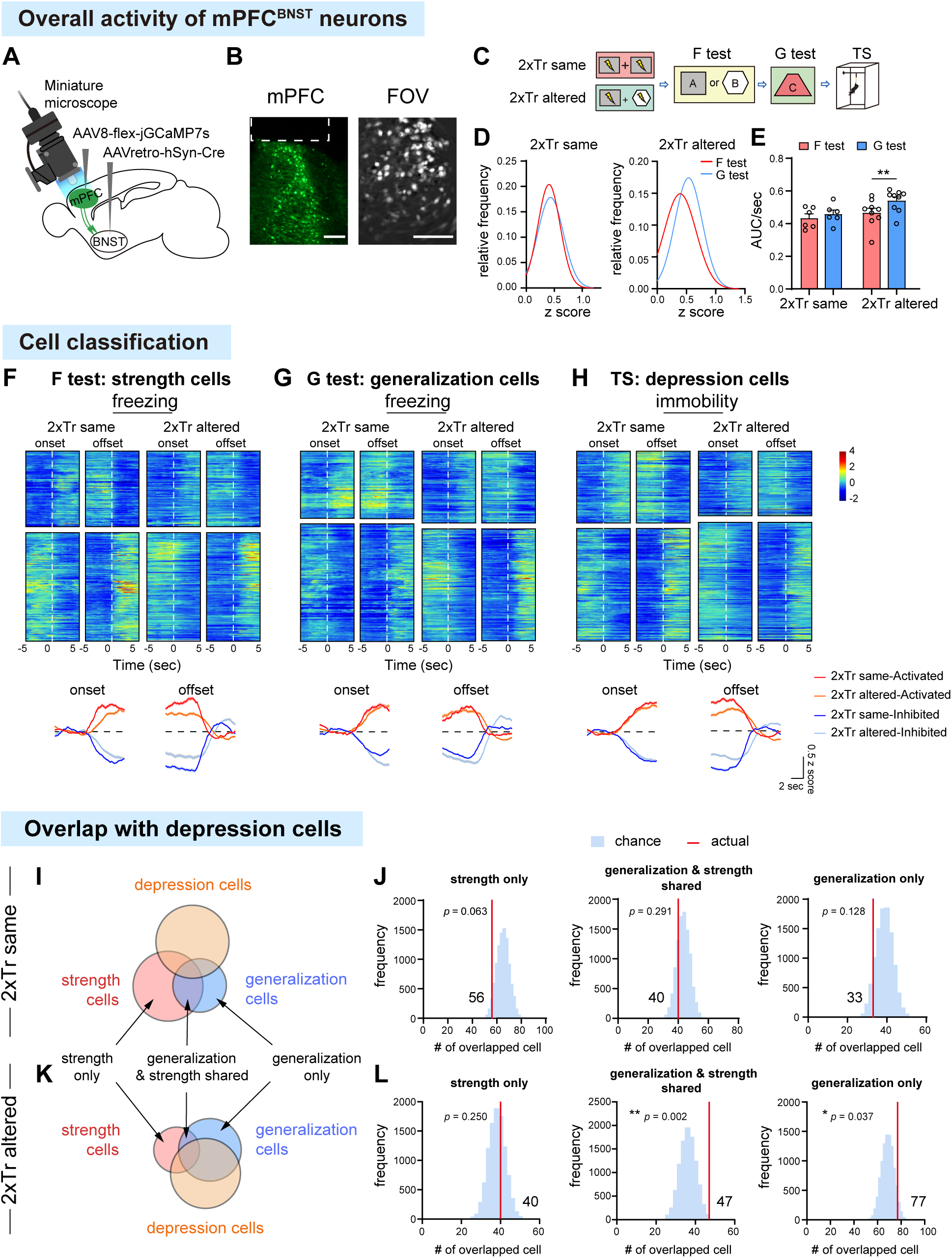
The mPFC^BNST^ neuronal ensemble encoding the generalization of negative memory, but not its strength, overlaps with that encoding depression-like behavior following variable negative experiences. (**A**) Schematic of AAV infusion, lens implantation and attachment of miniaturized microscope. (**B**) Representative images of jGCaMP7s expression (Scale bar, 300 µm) and a field of view (FOV, Scale bar, 150 µm) in the mPFC. (**C**) Schematic of two CFC trainings in the same (2xTr same) or different contexts (2xTr altered), fear memory strength test (F test), generalization test (G test) and tail suspension (TS) test. (**D, E**) Distribution (D) and quantification (E) of the overall calcium activity during memory tests after “2xTr same” (6 animals; 804 cells for F test, 842 cells for G test) or “2xTr altered” (9 animals; 810 cells for F test, 814 cells for G test). (**F-H**) Top: heatmaps of mean responses of activated and inhibited cells in response to freezing or immobility, aligned to the onset and offset of behaviors after “2xTr same” or “2xTr altered”. Bottom: averaged responses of the activated or inhibited cells. Lines indicate the mean value and shades indicate the s.e.m. (**I, K**) The percentages of “strength cells”, “generalization cells” and “depression cells” in the total number of recorded cells, and their overlaps. Schematic for defining “strength only”, “generalization only” (responding to freezing only in the F or G test) and “generalization & strength shared” (responding to freezing in both memory tests) cells. (**J, L**) Histograms indicating the distribution of the number of cells by chance (blue shade), compared with the observed number (red line), overlapping between “depression cells” and “strength/generalization cells” after “2xTr same” (J) or “2xTr altered” (L).

Analysis of Ca^2+^ activity during CFC training sessions revealed that a majority of visualized mPFC^BNST^ neurons were either activated (47-56%) or inhibited (33-37%) by foot-shocks (**Supplementary Fig. 4**). After the “2xTr altered” but not “2xTr same” paradigm, the mPFC^BNST^ neurons exhibited significantly higher overall activity during the memory generalization test compared to the memory strength test, suggesting that these neurons developed an increased response to context change following variable negative experiences (**Fig. 4D-E**).

We further aligned the Ca^2+^ activity of mPFC^BNST^ neurons to the onset or offset of freezing behavior during memory tests and immobility behavior during tail suspension, and then utilized the K-means clustering, an unsupervised machine learning algorithm, to categorize these neurons (**Supplementary Fig. 5**). We defined the mPFC^BNST^ neurons that exhibited activity changes in response to freezing during memory generalization test and strength test as “generalization cells” and “strength cells” respectively, and those that showed activity changes in response to immobility during tail suspension were referred to as “depression cells” (**Fig. 4F-H**). To determine overlaps between these cell populations, we conducted a permutation test and compared the actual number of overlapped cells to the null distribution. The results showed significant overlaps of the “depression cells” with the “generalization and strength shared cells”, as well as with those identified only as “generalization cells” following the “2xTr altered” paradigm, but not the “2xTr same” paradigm (**Fig. 4I-L**). Notably, no significant overlap was observed between “depression cells” with those solely identified as “strength cells” (**Fig. 4I-L**).

Instead, there was a trend indicating separation between these two neuronal ensembles after the “2xTr same” paradigm which increased memory strength (**Fig. 4J**).

Collectively, these findings suggest that as regards to neuronal encoding ensembles, variable negative experiences induce an increase in overall mPFC^BNST^ neuronal activity in response to context change, and a significant overlap of mPFC^BNST^ neuronal ensemble encoding depression-like behavior and negative memory generalization, but not with those encoding memory strength.

### Connectivity of the mPFC-to-BNST projections

To gain an understanding of the connectivity properties of the mPFC-to-BNST projections, we examined the distribution and cell-type of mPFC^BNST^ neurons. The results showed that within the mPFC subregions, a substantial proportion of the BNST-projecting neurons located in the deep layers of the infralimbic sub-division, and majority of these neurons were excitatory neurons (**Supplementary Fig. 6A-E**). Retrograde trans-monosynaptic tracing with recombinant rabies virus revealed that the mPFC^BNST^ neurons received monosynaptic inputs primarily from the ACC and the anterodorsal and mediodorsal thalamus (AD/MD), which are involved in information integration (**Supplementary Fig. 6F-H**)^32, 33^. In the BNST, the oval subregion received the highest proportion of mPFC projections, while other BNST subdivisions also received significant inputs from the mPFC. The BNST neurons targeted by the mPFC projections (^mPFC^BNST) were predominantly inhibitory neurons (**Supplementary Fig. 6I-N**). Anterograde trans-monosynaptic tracing with AAV1 showed that the ^mPFC^BNST neurons projected to various subcortical regions involved in emotion processing, including VTA and LHb (**Supplementary Fig. 6O-P**).

### Activation of the mPFC-to-BNST projections promotes depression-like behaviors

We next asked after negative experiences, whether manipulations of the mPFC-to-BNST projection, which supported generalization of negative memories but not memory strengthening, could lead to changes in depression-like behaviors. We specifically expressed hM4Di in the mPFC^BNST^ neurons and trained the mice with the “2xTr altered” paradigm (**Fig. 5A**). Suppression of the mPFC^BNST^ neuronal activity ameliorated depression-like behaviors, as indicated by a decrease in latency to eat in the novelty-suppressed feeding test, an increase in grooming duration in the splash test, a decrease in immobility in the tail suspension test and an increase in sociability. These effects were observed both during the first week and after two weeks following the “2xTr altered” paradigm (**Fig. 5B-E, Supplementary Fig. 7A-E**).

**Fig. 5.**
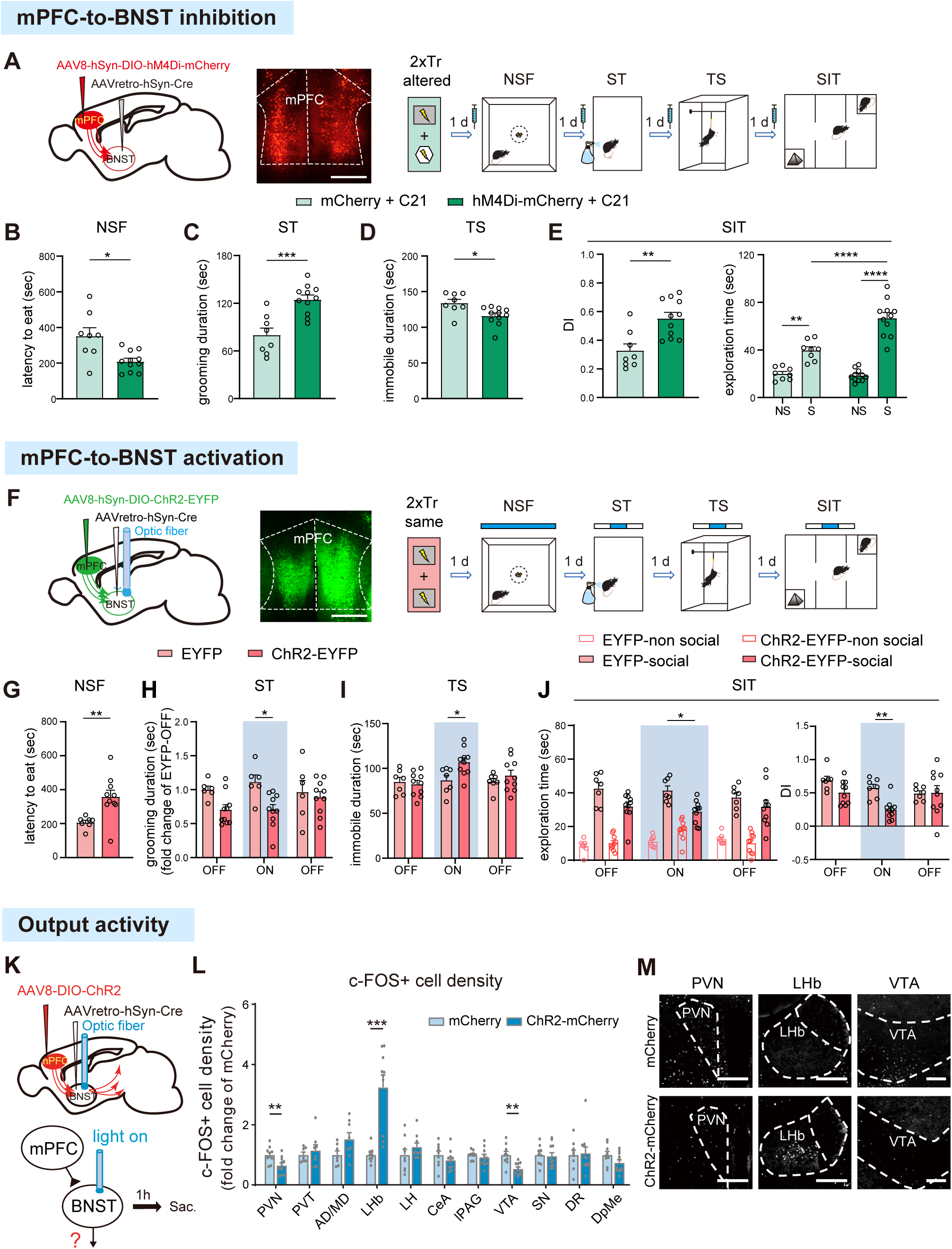
Activation of the mPFC-to-BNST projections promotes depression-like behaviors. (**A, F**) Schematics of AAV infusion, representative images of mCherry or EYFP expression in the mPFC and experimental timelines. Scale bar, 500μm. (**B, G**) Quantification of the latency to eat in novelty-suppressed feeding (NSF) test. (**C, H**) Quantification of the grooming duration in splash test (ST). (**D, I**) Quantification of the immobile time during tail suspension (TS) test. (**E, J**) Quantification of the exploration time and differential index (DI) in social interaction test (SIT). B-E, changes in behaviors by chemogenetic inhibition of the mPFC-to-BNST projections after “2xTr altered”. n = 8-11 per group. G-J, changes in behaviors by optogenetic activation of the mPFC-to-BNST projections after “2xTr same”. n = 6-10 per group. (**K**) Schematic of AAV injection, optogenetic activation and sample collection (sac). (**L**) Fold changes in the density of c-Fos+ cells across different downstream regions of BNST in response to optogenetic activation of the mPFC-to-BNST projection. n = 7-11 per group. (**M**) Representative images of c-FOS staining in the periventricular nucleus of hypothalamus (PVN), the lateral habenula (LHb) and the ventral tegmental area (VTA) after optogenetic activation of the mPFC-to-BNST project. Scale bar: 200 μm. Data are presented as mean ± s.e.m. and analyzed by *t* tests and two-way ANOVA followed by Sidak’s *post hoc* test. * *p* < 0.05, ** *p* < 0.01, *** *p* < 0.001, **** *p* < 0.001.

In the reverse fashion, we expressed ChR2 in the mPFC^BNST^ neurons and implanted fibers in the BNST for optogenetic activation of the mPFC-to-BNST projection (**Fig. 5F, Supplementary Fig. 3C**). After the “2xTr same” paradigm, activation of the mPFC-to-BNST projection enhanced depression-like behaviors both during the first week and after two weeks following the last training (**Fig. 5G-J, Supplementary Fig. 7F-J**). Consistent with the behavioral phenotypes, optogenetic activation of the mPFC-to-BNST projection resulted in activation of the lateral habenula (LHb) and decreased activities in the ventral tegmental area (VTA), resembling what is commonly observed in depression patients and animal models (**Fig. 5K-M**)^34, 35^.

Collectively, these findings demonstrate that at the neural circuit level, activity of the mPFC-to-BNST projection promotes both generalization of negative memories and depression-like behaviors, but not memory strengthening.

### Activation of the consolidation mechanism specific to memory generalization elicits depression-like behaviors

To further investigate the hypothesis that negative memory generalization, as opposed to memory strengthening, is sufficient to trigger depression, we postulated that activation of the consolidation process specific to memory generalization, rather than strengthening, following negative experiences would elicit depression-like behaviors. The consolidation process necessary for the formation of long-term memory involves *de novo* gene expression^36, 37^. To identify how consolidation mechanisms of memory generalization differ from memory strengthening, we utilized the Retro-TRAP (translating ribosome affinity purification) method to capture transcriptional changes in mPFC^BNST^ neurons following training^38, 39^. We selectively expressed NBL10 (anti-GFP Nanobody-Rpl10a fusion) in the mPFC^BNST^ neurons, and subjected the mice to different patterns of training. One hour after the last training, the mPFC was collected for immunoprecipitation of NBL10-tagged ribosomes, from which the attached RNAs were purified and sequenced (**Fig. 6A**). We observed an enrichment of excitatory neuronal markers and a depletion of inhibitory neuronal and glial markers in the TRAP samples compared to the total inputs, consistent with the predominately excitatory nature of the mPFC^BNST^ neurons (**Fig. 6B**)^40,41^.

**Fig. 6.**
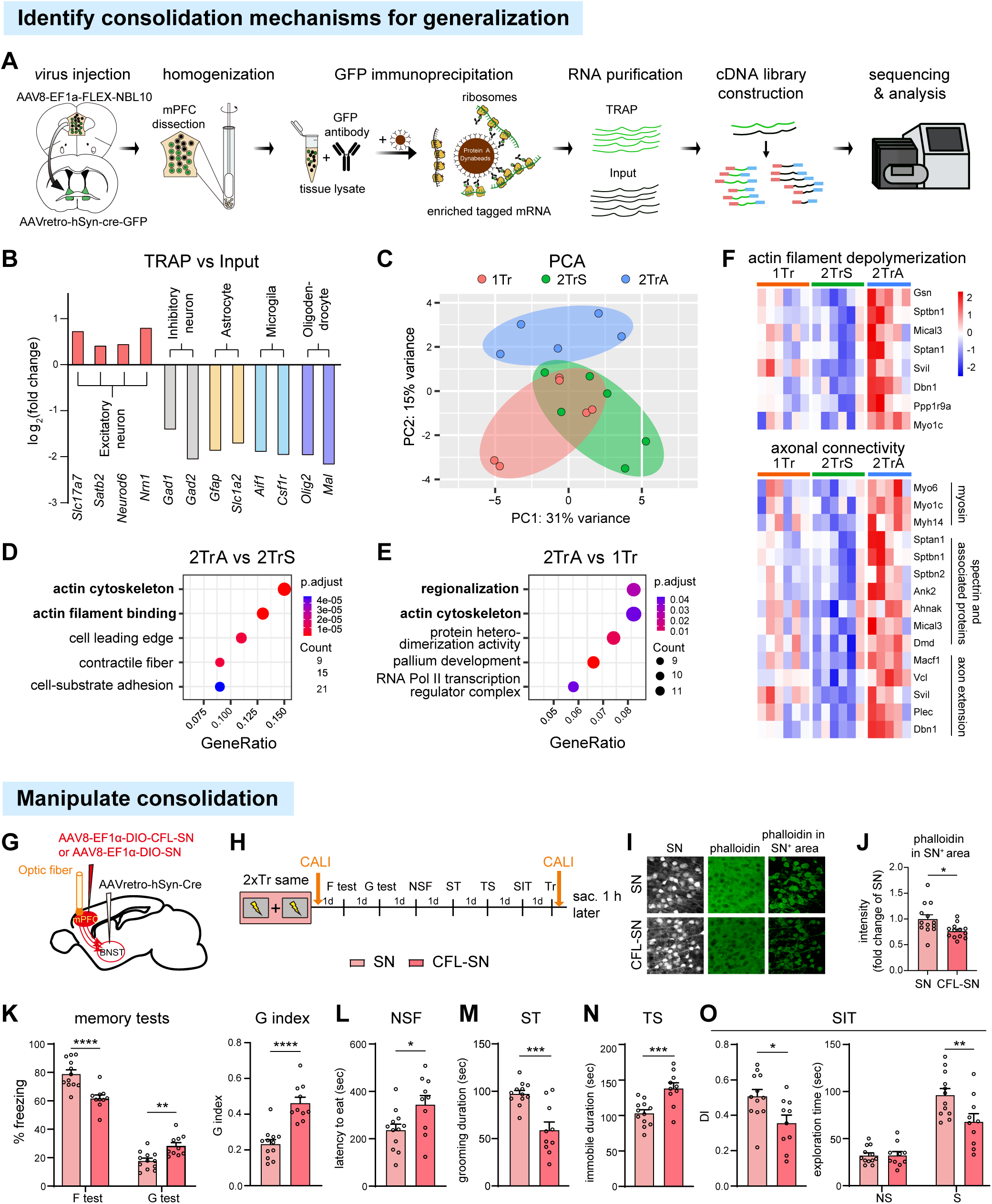
Activation of the consolidation mechanism specific to memory generalization elicits depression-like behaviors. (**A**) Schematic of the Retro-TRAP procedure. (**B**) Changes in expression of neuronal and glia marker genes, comparing translating ribosome affinity purified (TRAP) versus input RNA. (**C**) Principal Component Analysis (PCA) plot of TRAP samples. 1Tr: a single CFC training in either context A or B; 2TrS: two CFC trainings in the same context; 2TrA: two CFC trainings in altered contexts. (**D-E**) Top 10 enriched Gene Ontology of DEGs comparing the 2TrA versus 2TrS (D) and 2TrA versus 1Tr (E). (**F**) Heatmaps showing the relative expression of DEGs related to actin cytoskeleton across groups. (**G**) Schematic of AAV infusion and optic fiber implantation for inducing actin remodeling in the mPFC^BNST^ neurons. (**H**) Schematic of behavior paradigm and chromophore-assisted light inactivation (CALI) application. (**I-J**) Representative images and quantification of phalloidin staining in the experimental group expressing cofilin fused to SuperNova (CFL-SN) and the control mice expressing SuperNova (SN) only. Scale bar, 50 μm. n = 9 per group. (**K**) Quantification of the percentage of time spent in freezing during the memory strength test (F test) and the generalization test (G test), and the G index. (**L-O**) Quantification of the latency to eat in novelty-suppressed feeding (NSF) test (L), the grooming duration in splash test (ST) (M), the immobile time during tail suspension (TS) test (N), the exploration time and differential index (DI) in social interaction test (SIT) (O). n = 10-12 per group. Data are presented as mean ± s.e.m. and analyzed by *t* tests and two-way ANOVA followed by Sidak’s *post hoc* test. * *p* < 0.05, ** *p* < 0.01, *** *p* < 0.001.

We further analyzed the TRAP RNAs by comparing the “2xTr same” with the “1xTr” groups to identify consolidation mechanisms for memory strengthening, and comparing the “2xTr altered” with the “1xTr” and “2xTr same” groups to identify mechanisms for memory generalization. Principal Component Analysis revealed that the “2xTr altered” group was distinct from the “1xTr” and “2xTr same” groups, whereas the latter two showed a partial overlap (**Fig. 6C**). Gene Ontology analysis of the differentially expressed genes (DEGs) identified enrichment in transcriptional changes related to actin cytoskeleton in the “2xTr altered” group compared to the other two groups (**Fig. 6D-E**), whereas changes in the “2xTr same” versus “1xTr” comparison were enriched for extracellular matrix and protein serine kinase activity, suggesting distinct transcriptional alterations associated with memory strengthening versus generalization. Further analysis of the DEGs related to actin cytoskeleton showed upregulation of genes involved in actin depolymerization and axonal alterations (**Fig. 6F**)^42–47^.

To validate the involvement of actin remodeling during memory consolidation for generalization and to explore its impact on the onset of depression-like behaviors, chromophore-assisted light inactivation (CALI) was employed to induce acute actin destabilization in mPFC^BNST^ neurons during the memory consolidation phase (**Fig. 6G**)^48–50^. Cofilin fused with the photosensitizer SuperNova was expressed in mPFC^BNST^ neurons, and 593 nm illumination was used to generate reactive oxygen species by SuperNova, leading to cofilin inactivation and subsequent actin cytoskeleton destabilization^48–50^. Mice were trained with the “2xTr same” paradigm, which typically results in memory strengthening without generalization. Light stimulation was applied to trigger actin destabilization in the mPFC^BNST^ neurons 2 min after the second training (**Fig. 6H-J, Supplementary Fig. 3E**). This manipulation altered the paradigm’s outcome, leading to memory generalization while preventing memory strengthening (**Fig. 6K**). Importantly, it also rendered the “2xTr same” paradigm to induce depression-like behaviors (**Fig. 6L-O**).

Notably, the CALI manipulation occurred during the post-training memory consolidation phase, thereby eliminating any potential effect on the perception of unpredictability and the associated high stress levels during training. Therefore, our findings underscore the crucial role of memory components in depression. Furthermore, these data support the hypothesis that enhanced memory generalization is a crucial factor coupled with the onset of depression, challenging the traditional notion that the strengthening of negative memories contributes to depression.

## DISCUSSION

Negative experiences, recognized as an external factor contributing to the onset of depression, encompass two elements: stress during the events and lasting memories thereafter. Diverging from conventional depression research, which predominantly emphasizes the stress component, our study delves into the pivotal role of memory in the etiology of depression. By introducing behavioral paradigms that differentiate between memory strengthening and generalization, our study provides empirical evidence across four different levels of analysis (behavioral, neural circuit, neuronal ensemble activity, and molecular), which collectively indicates that the generalization of negative memories, rather than their strengthening, essentially drives depression-like behaviors.

Memories of past experiences are integral to human mental well-being^12, 16^. While the exact cause of depression remains unclear, cognitive theories posit that maladaptive negative memories contribute to negative thought patterns, thereby influencing the onset and maintenance of depression. Depressed individuals preferentially recall and intensify negative personal memories, fostering rumination and subsequent overgeneralization of these memories^4, 5, 8^. However, experimental evidence clarifying the relative importance of memory reinforcement or generalization in fueling depression has been lacking, partly due to the intricate interplay between these two memory components. To address this gap, we developed new mouse behavioral paradigms revealing that repeated negative experiences in the same context strengthen memories, whereas negative experiences in varying contexts enhance generalization. These paradigms not only establish a framework for exploring the mechanistic link between memories and neuropsychiatric disorders, but also provide behavioral evidence suggesting that the overgeneralization of negative memories, rather than their strengthening, is intimately tied to depression-like behaviors.

Our findings resonate with clinical and animal studies reporting that exposure to diverse negative experiences, as opposed to repeated exposure to the same stressor, escalates the risk of depression^21, 51^. Importantly, our study offers further insights into how memories of unpredictable negative experiences differ from those of predicable ones and how such differences influence depression-like behaviors. We demonstrated that post-training manipulations of a neural circuit promoting memory generalization, which did not affect the perception of unpredictability or the associated high stress responses during training, could alter depression-like behaviors. These results underscore the critical role of negative memories beyond stress in depression. Our findings suggest that unpredictable stressful experiences exacerbate the overgeneralization of aversive memories, rather than intensifying their strength, ultimately leading to depression-like behaviors. This addresses a knowledge gap in the field. Based on these findings, we propose that cognitive therapies focusing on reducing overgeneralization and improving the accuracy of past negative memories may be more beneficial for individuals with depression, compared to efforts to extinguish or alter the emotional valence of memories. Additionally, assessing the precision of negative memories could serve as a cognitive marker for evaluating the risk and severity of depression in individuals and their response to therapeutic interventions.

The current research not only underscores the significant contribution of negative memory generalization to depression, but also advocates for deeper mechanistic investigation of memory quality. While prior research has made strides in unraveling the regulatory mechanisms of memory strength^37, 52^, the mechanistic understanding of memory generalization remains limited. From a brain network perspective, it is unclear whether distinct brain regions and circuit connections mediate these two components of memory. Our study, leveraging c-FOS mapping, functional connectivity analysis, circuit tracing, and manipulations, uncovered that the BNST serves as the central hub for the brain network activated by variable negative experiences that promote memory generalization, whereas the CeA functions as the core hub for networks activated by experiences that generate precise memories. These findings indicate that memory generalization and strength rely on distinct neural circuits. Furthermore, while the amygdala has garnered significant attention in fear memory research^53^, our results underscore the BNST’s pivotal role for the expression of conditioned fear memory following variable negative experiences. We further found that activation of mPFC-to-BNST projection promotes memory generalization and depression-like behaviors, but not memory strengthening. Employing single-cell imaging of calcium activity, we observed that training in different contexts increased the overall responsiveness of mPFC^BNST^ neurons to contextual changes, resulting in significant overlap in the mPFC^BNST^ neuronal ensembles encoding depression-like behaviors and generalization, but not memory strength. Given the mPFC’s involvement in schema memory^54^ and the BNST’s role as a relay station regulating the hypothalamic-pituitary-adrenal axis^25^, the mPFC-to-BNST projection emerges as a critical circuit mediating the influence of schema memory on stress response, ultimately leading to maladaptive behaviors. This projection represents a potential target for neuromodulation therapies aimed at reducing memory generalization and depression^55, 56^.

Our findings, together with other studies indicating complex regulation of memory generalization by BNST, mPFC and VTA (which is downstream of the mPFC-to-BNST projection), collectively advocate for more comprehensive investigation into the regulation and connectivity of mPFC-to-BNST circuit during various stages of memory generalization process ^57–60^. It is also important to note that the mPFC contains subdivisions, with functional differences in fear regulation and depression ^29, 61^. A limitation of our study is lacking the functional differentiation between the mPFC subdivisions. In examining the mPFC-to-BNST projection, our AAV injections were primarily targeted at both the PL and IL subdivisions of the mPFC, with a substantial proportion of BNST-projecting neurons identified within IL. Further research is needed to elucidate the distinct roles of different mPFC subdivision in linking memory generalization to depression.

Contrary to traditional depression research often using chronic stress paradigms in animal studies^9, 10, 62^, our study demonstrates that two negative experiences in variable contexts can elicit depression-like behaviors persisting for at least two weeks. This aligns with clinical observations linking acute stressful events to the onset or recurrence of major depression episodes^63, 64^. Using Retro-TRAP, we found that memory enhancement and generalization may be governed by distinct molecular pathways. By temporally and cell-type specific inducing actin remodeling in the mPFC^BNST^ neurons during memory consolidation, we observed compromised memory quality, characterized by reduced memory strength but increased memory generalization. Furthermore, our manipulation, without affecting stress level during training, was sufficient to induce depression-like behaviors. These results imply that not only is the strengthening of negative memories less crucial than previously thought^12, 14–16^, but also the degree of their generalization is a far more significant factor in driving depression-like behaviors. Additionally, our findings also highlight actin remodeling as a potential molecular target for developing effective treatments aimed at reducing excessive generalization of negative memories in depression.

In summary, our findings underscore the need of addressing the overgeneralization of negative memories in the context of depression interventions. Early efforts in this direction have demonstrated promising outcomes^17^. Future research should prioritize the development of enhanced psychotherapy protocols, potentially combined with neuromodulation strategies, to achieve more effective memory specificity training. By identifying the key underlying neural circuits and molecular mechanisms, our findings lay the groundwork for the development of novel interventions for the prevention and treatment of depression. Given that the overgeneralization of negative memories is a prevalent cognitive symptom across a spectrum of stress-related neuropsychiatric disorders^65, 66^, this study also offers insights into the mechanisms linking maladaptive memories with these conditions.

## ACKNOWLEDGMENTS

We thank Dr. Thomas J. Carew at New York University and Dr. Xueyi Shen at University of Edinburgh for helpful comments on the manuscript. We thank the animal facility and the core facility of the Zhongshan School of Medicine, Sun Yat-sen University. We thank all the lab members for discussion and technical assistance during the execution of this project. This work is supported by grants from the STI2030-Major Projects (No. 2021ZD0202000 to XY and YNZ), National Natural Science Foundation of China (No. 32271068 and No. 81873797 to XY, No. 81972967 to WJL, No. 81971270 to YNZ), Guangzhou Science and Technology Program key projects (No. 202007030001 to XY and WJL), Guangdong Science and Technology Department (No. 2020B1212060018 and 2020B1212030004 to WJL), the Science and Technology Planning Project of Guangdong Province (No. 2023B1212060018 to XY and WJL) and Guangdong Project (No. 2019QN01Y202 to XY).

