## Supplemental Materials for "Generalization but not strengthening of negative memories drives the development of depression-like behaviors"

Running title: negative memory generalization drives depression

Xin Cheng, Ph.D.^1†^, Yubo Hu, Ph.D.^1,2†^, Yan Zhao, Ph.D.^1†^, Panwu Zhao, M.S.^1^, Xiaomeng Bai, M.S.^1^, Yi Chen, B.S.^1^, Deshan Kong^1^, Shuyu Zheng, B.S.^1^, Yuena Zheng, M.S.^1^, Yumeng Wang, B.S.^1^, Yanni Zeng, Ph.D.^1^*, Wei-Jye Lin, Ph.D.^3,4,5^*, Xiaojing Ye, Ph.D.^1^*

**Supplementary materials and Methods**

**Animals**

8 to 12-week-old male and female C57BL/6J mice at the start of the experiments were used in the studies. The mice were obtained from the Institute of Experimental Animals of Sun Yat-sen University (Guangzhou, China), Guangdong Medical Laboratory Animal Center (Guangzhou, China) or Zhuhai Bai Shi Tong Animal Center (Zhuhai, China). The animals were housed in groups of 4-5 at 23 ± 2°C and 50~60% humidity on a 12 h light/dark cycle in a specific pathogen free (SPF) facility with water and food available ad libitum. The animals were allowed to accustom to the animal facility for at least one week after transportation, before initiation of any experimental procedures. All behavioral testing was performed during light cycle, and animals were randomly assigned to different experimental groups. All animal studies were reviewed and approved by the Institutional Animal Care and Use Committee of Sun Yat-sen University under the approval number SYSU-IACUC-2018-000065 and SYSU-IACUC-2022-000492.

**CFC training and testing**

Mice were handled for 2-3 min/day for 4 days before training. Four contexts were used in the study: context A was a rectangular fear conditioning chamber with clear walls, illuminated with 90 lux natural white light. The floor was made of stainless-steel rods connected to a shock delivery apparatus (Med Associates, St. Albans, Vermont, USA). The chamber was placed in an acoustically-insulated cabinet and cleaned with 4% acetic acid between animals. Context B was a chamber modified with red-and-white striped walls and illuminated with 90 lux red light. The floor was made of stainless-steel rods connected to a shock delivery apparatus (Shanghai Jiliang Software Technology Co., Ltd., China). The chamber was placed in a different acoustically-insulated cabinet and cleaned with 75% alcohol between animals. Context C was a triangle clear acrylic chamber placed in the acoustically-insulated cabinet as used for context A or B. The chamber was illuminated with 60 lux red light and cleaned with 2% ammonia between animals. Context A’ used the same chamber as context A, modified with a curved white wall insert and a white acrylic floor, and illuminated with 60 lux natural white light in the acoustically-insulated cabinet as used for context A.

CFC consisted of a 3-min habituation to the context A or B followed by three 2-s foot shocks (0.75 mA) with an inter-shock interval of 60 s (*62*). After the third shock, animals were kept in the context for an additional 60 s. On the following day, the animals in the “two train” groups underwent another CFC training in the same context or a new context. The next two days, mice were re-exposed to the context A or B for fear memory strength tests (F test), and a modified context (context C or A’) for memory generalization tests (G test), with an interval of 24 hours. The temporal order of memory strength and generalization tests was counterbalanced between animals. During testing, freezing behavior was videotaped. Analysis of the freezing responses was conducted offline by an examiner blinded to the experimental groups. Freezing was defined as the absence of any visible movements other than those necessary for respiration. The percentage of time spent in freezing during the 3 min tests was measured.

We trained mice with one of the three paradigms: (1) “1xTr”: mice received single CFC training in either context A or B, with counterbalanced selection between animals. (2) “2xTr same”: mice received two CFC training sessions in the same context (A or B, counterbalanced between animals), spaced by 24 h. For both “1xTr” and “2xTr same” groups, F test was conducted in the same context as training and G test were conducted in the context C, except for the similar context experiment (context A’) as specified in the main text. (3) “2xTr altered”: mice received two CFC training sessions in different contexts (A and B, the temporal order of which is counterbalanced between animals), spaced by 24 h. F test was conducted in either context A or B, with counterbalanced selection between animals. G test was conducted in context C.

**Stereotaxic surgery**

Mice were anesthetized with 1.5% isoflurane at an oxygen flow rate of 0.4 L/min, and head-fixed on a stereotactic frame (RWD Life Science, Shenzhen, China) equipped with a heating pad to maintain body temperature at 37°C. Eyes of mice were lubricated with Vaseline mixed with saline. After shaving the fur and sterilizing the incision site, an incision was made to expose the skull. Holes were drilled in the skull. Virus was injected by a glass pipette at 20-50 nl/min with a micro-syringe pump (Nanoliter2020, WPI, Sarasota, FL, USA). After injection, the pipette was left at the injection site for an additional 5 min before slow withdrawal to allow diffusion of the virus and minimize backflow. Adeno-associated virus (AAV) and rabies virus (RV) used in the study were purchased from Obio Technology (Shanghai, China), BrainVTA (Wuhan, China) and BrainCase (Shenzhen, China).

For retrograde tracing of upstream neurons projecting to the BNST, 200 nl 0.5% Cholera Toxin Subunit B conjugated with Alexa Fluor 555 (CTB-AF555) was injected per side bilaterally at the BNST (AP: +0.3 mm, ML: ±1.0 mm, DV: −4.35 mm). For optogenetic activation of mPFC or vSUB projections to the BNST, mice were injected with 100 nl of AAV8-CaMK2a-hChR2(E13A)-mCherry (1.53 ×10^13^ vg/ml) or AAV8-hSyn-DIO-hChR2(E13A)-mCherry (2.36×10^13^ vg/ml) into the mPFC (AP: +1.8 mm, ML: ±0.3 mm, DV: −2.45 mm) or vSUB (AP: -3.8 mm, ML: ±3.3 mm, DV: −4.5 mm) and then implanted unilaterally with optic fibers (200-µm core diameter, Inper) above the BNST. AAV8-CaMK2a-mCherry (1.27×10^13^ vg/ml) or AAV8-hSyn-DIO-mCherry (2.36×10^13^ vg/ml) was used as the controls. For chemogenetic inhibition of the mPFC, ACC and LS projections to the BNST, mice were bilaterally injected with 200 nl per side of AAVretro-hSyn-EGFP-2A-CRE (1.63×10^13^ vg/ml) into the BNST and 100 nl per side of AAV8-hSyn-DIO-hM4D(Gi)-mCherry (5.23×10^12^ vg/ml) into the mPFC, ACC (AP：+1.0 mm，ML：±0.3 mm，DV：-1.6 mm) or LS (AP：+0.8 mm，ML：±0.4 mm，DV：-3.5 mm). AAV8-hSyn-DIO-mCherry (5.00×10^12^ vg/ml) was used as the controls. For CALI-induced actin remodeling in the mPFC^BNST^ neurons, mice were injected with 100 nl per side of AAV8-EF1a-DIO-CFL-SN (5.95 ×10^12^ vg/ml) or AAV8-EF1a-DIO-SN (5.78 ×10^12^ vg/ml) as a control into the mPFC and 200 nl per side of AAVretro-hSyn-CRE (6.25×10^11^ vg/ml) into the BNST. Optic fibers were implanted bilaterally above the mPFC.

For labelling different BNST neurons downstream of the mPFC, 200 nl per side AAV8-hSyn-DIO-mCherry (7.00×10^12^ vg/ml) was injected bilaterally into the BNST and 100 nl AAV1-hSyn-EGFP-2A-CRE (3.72×10^13^ vg/ml) per side was injected into the mPFC, as high titer AAV1 can be anterogradely and trans-synaptically transported (*63*). For tracing upstream inputs of the mPFC^BNST^ neurons, 100 nl per side helper viruses consisted of 1:1:1 mixture of AAV9-EF1α-DIO-H2B-EGFP-T2A-TVA (2.10×10^12^ vg/ml), AAV8-EF1α-DIO-oRVG (2.08×10^12^ vg/ml) and AAV8-CMV-CRE (9.39×10^13^ vg/ml) were injected bilaterally into the mPFC. After 3 weeks, 200 nl per side RV-ENVA-△G-dsRed (2.00×10^8^ IFU/ml) was injected bilaterally into the BNST. The presence of these helper viruses enabled the RV to spread retrogradely across mono-synapses (*64*).

**Depression-like behavioral tests**

In the study, we employed a battery of four behavioral assays, which are well-established in preclinical research for probing depression-like phenotypes in rodents. Their combined use in our study provides a comprehensive assessment of depression-like behaviors, addressing the multifaceted nature of depression. Furthermore, our study not only measured behavioral changes starting one day after the last training but also examined their temporal stability for at least two weeks after the last training session.

Depression-like behaviors were assessed starting at 1 day and 2 weeks after the last training with a 24 h-gap between tests, as previously described (*26*). Mice were brought to the procedure room 2 h before each behavioral test for acclimation. The behavioral apparatus was cleaned with 75% ethanol between animals. The order of the behavioral tests was specified in the experimental timeline in the related figures.

*Novelty suppressed feeding:* After food deprivation for 24 h, mice were placed into one of the corners of a white acrylic box (40 × 40 × 40 cm) with one food pellet placed in the center. The box was illuminated under red-light (80–100 lux) in a sound-insulated chamber. The video was recorded was DigBeh system (Shanghai Jiliang Software Technology Co., Ltd., China). The latency to begin food consumption in a 10-min test was recorded by a researcher blinded to the experimental groups.

*Splash test:* The test was carried out in a mouse cage illuminated under red-light (80–100 lux) in a sound-insulated chamber. The back fur of mouse was sprayed 3 times with 10% sucrose solution and the mouse was videotaped for 5min. The time spent grooming was analyzed by a skillful researcher blinded to the experimental groups.

*Social interaction test:* A three-chamber social interaction test was performed in an arena (60 × 40 × 20 cm) with three separate chambers illuminated under 80-100 lux red light. A social mouse (sex and age matched) unfamiliar to the test mouse was habituated in a cup placed in the arena for 10 mins one day before the test. On the test day, the test mouse was allowed to explore freely for 10 min the arena with two empty cups placed in the corners of side chambers. Afterward, the unfamiliar social mouse was placed into one of the two cups, while the other contained a toy mouse. The test mouse was placed back into the middle chamber and videotaped for 10 mins. The time spent investigating the social mouse or toy mouse was measured offline by a researcher blinded to the experiment groups. The social preference differential index (DI) was calculated by the following formula: (investigation time of a social mouse – investigation time of toy mouse) / (total investigation time of the social mouse and the toy mouse).

*Tail suspension:* Mice were suspended by their tails from a metal rod fixed 50 cm above the surface of the four-walled rectangular compartments under white light (100-120 lux). The tip of tail was passed through a 1 ml pipette tip and then fixed using adhesive paper tape. The test mouse was suspended by the tail for 6 min and videotaped. The immobile time was measured offline for the last 4 min by a researcher blinded to the experiment groups. The mice were considered as immobile when their four limbs stopped struggling, without substantial body movement.

**Optogenetic, chemogenetic and CALI manipulations**

For optogenetic experiments, a 473-nm blue laser diode (Thinker Tech Nanjing Bioscience Inc., China) was connected to the ferrule of the optic fiber implanted on mouse head. Laser power was measured at the fiber tip before each experiment and adjusted to ~3 mW. Each trial was consisted of a 20 Hz pulse train (10 ms per pulse) lasting for 120 sec. Four weeks after the virus injection and fiber implantation, mice underwent behavioral tests. Freezing time was calculated as the averaged time spent in freezing across light ON or light OFF epochs of the session. The photo-stimulation protocol was chosen based on a previous study (6*5*). For behavioral testing with chemogenetic manipulations, clozapine (Sigma-Aldrich, 0.1 mg/kg) or Compound 21 (C21, Tocris Bioscience, 1 mg/kg) was dissolved in 0.9% saline and injected intraperitoneally 1 h before testing. For CALI-induced actin cytoskeleton remodeling in the mPFC^BNST^ neurons, a 593-nm laser diode (Thinker Tech Nanjing Bioscience Inc., China) was connected to the ferrule of the optic fiber implanted on mouse head. Laser power was measured at the fiber tip before each experiment and adjusted to ~3 mW. Four weeks after the virus injection and fiber implantation, mice were trained with two CFC trainings in the same context, separated by 24 h. Two minutes after the second training, the mice received the 593-nm laser illumination through the optic fiber for 1 min (*48*). Behavioral testing of memory performance and depression-like behaviors were conducted starting 24 h after the last training in the absence of the laser light.

**Immunofluorescence staining and image analysis**

Mice were perfused transcardially with 1×PBS for 1 min and 4.0% paraformaldehyde in 1×PBS (pH 7.5) for 5 min under deep anesthesia. Brains were removed, post-fixed (4% paraformaldehyde in 1xPBS overnight, 4℃) and cryoprotected (30% sucrose, 1×PBS, 4 ℃, over 48 h). 40 μm coronal sections were collected with a cryostat (Leica, Germany). For c-FOS staining, mice were sacrificed 60 min after behavioral testing. For c-FOS staining upon hM4Di-mediated inactivation or ChR2-mediated activation, animals were sacrificed 60 min after drug injection or light application. Free-floating sections were incubated in a blocking buffer (1×PBS with 1% bovine serum, 5% normal goat serum, 0.4% Triton X-100) at room temperature for 2 h, followed by incubation with primary antibodies in the blocking buffer (1% BSA, 1×PBS, 0.4% Triton X-100) over 40 h at 4 °C under constant shaking. Sections were washed extensively with 1×PBS with 0.4% TritonX-100 and then exposed to secondary antibodies (Life Technologies) in the blocking buffer at room temperature for 2 h, after which the sections were washed again, stained with DAPI (5 µg/mL, Cell Signaling Technology), and mounted in anti-fade reagents on the glass slides. The primary antibodies used in this study include anti-GABA (1:5000, Millipore, MAB1572), anti-CAMKII (1:1000, Millipore, 05-532), anti-c-FOS (1:750, Cell Signaling Technology, 2250). F-actin was stained with Alexa Fluor 647-phalloidin (1:1000, Thermo, A22287). Images were acquired by the Nikon Eclipse Ni-U epi-fluorescent microscope (Tokyo, Japan) on a motorized stage, and automatically stitched, or Zeiss LSM800 or LSM780 confocal microscope (Jena, Germany). The images were then analyzed by Fiji ImageJ (National Institutes of Health, USA).

**Network analysis of c-FOS expression across multiple brain regions**

The density of c-FOS positive cells was calculated for each brain region by ImageJ (National Institutes of Health, USA). Regions-of-Interest (ROIs) were manually drawn according to the Allen Mouse Brain Atlas. c-FOS positive cells were automatically counted using a custom-written macro with the same parameters used in thresholding and analyzing particles for all the images. A total of 30 brain regions of interest were analyzed in this study (see Table S1 for a complete list). Pearson’s correlations of interregional c-FOS expression were analyzed, and correlations at a significance level of *p* < 0.05 were used to generate networks using igraph (version 1.0.1) in R (version 3.2.2). The nodes in the networks represent each brain region, whereas the edges represent significant functional interactions (*21*). We then calculated the global efficiency of the networks using the braingraph (version 0.62.0) packages in R with the following equation:

$$Global efficiency=1/N(N-1)\sum_{(i\neq j\in G)} (\frac{1}{{d\_}_{ij}})$$

N refers to the number of nodes in the network,$d\_ij$ refers the distance of the node_i_ and node_j_, G refers to the network G.

To evaluate the contribution of each brain region in the network, each node was isolated one-by-one *in silico*, and the resulting changes in the network global efficiency were calculated and ranked (*21-23*).

**Single-cell calcium imaging in free-moving animals**

*Surgery:* For endoscopic single-cell calcium imaging of mPFC^BNST^ projection neurons, mice were injected unilaterally with 200 nl of AAVretro-hSyn-SV40 NLS-CRE (1.00×10^13^ vg/ml) into the BNST and 100 nl AAV9-FLEX-GCaMP7s (2.76×10^12^ vg/ml) into the mPFC. Three weeks after virus injection, a gradient refractive index (GRIN) lens (ProView TM Lens probe, 1.0 mm diameter, ~4.0mm length, Inscopix) was implanted into the mPFC during a second surgery. Under anesthesia, an incision was made to expose the skull. A hole was drilled in the skull above the mPFC (AP: +1.8 mm, ML: ±0.3 mm) and gently expanded to allow the lens to move through. The tissue 1 mm above the mPFC was aspirated using a vacuum pump. Then the lens was descended until an adequate plane of cell bodies and/or diffuse fluorescence was visible (DV: -2.5 to -3.0 mm as verified by post-experiment histological analyses) with the miniaturized microscope (nVista 3.0, Inscopix). The lens was implanted with a lateral angle of 15^o^ to avoid damaging major blood vessels at the midline. Extra space between the lens and the skull was filled with silicone adhesive (Kwik-sil, World Precision Instruments), and the lens was fixed to the skull with dental acrylic (C&B Superbond, NISSIN). Two weeks later, a third surgery was performed. A magnetic baseplate was lowered to the lens until a clear focal plane was visible with the miniaturized microscope. The baseplate was secured above the lens with dental acrylic.

*Image acquisition:* Ca^2+^ imaging in freely moving mice during different behavioral tests was performed via the miniaturized microscope (nVista 3.0, Inscopix) (*66*). Mice were habituated to the microscope attachment procedure for at least 4 consecutive days before experiments. Calcium images were acquired using the nVista acquisition software at 1280×800 pixels, with an exposure time of 50 msec and at a frame rate of 20 Hz. Light-emitting diode (LED) power setting was optimized for each animal in a range of 20 to 80% intensity (0.4 to 1.6 mW, 473 nm), with gain of 3 to 6, depending on the dynamic range of pixel values in the field of view.

*Image processing:* Raw movies were pre-processed by resizing the screen border and down-sampling to the 10 Hz. Motion artifacts were corrected and a mean filter was applied to the movies. A single frame average projection was created and used as the background fluorescence (F_0_) to calculate the normalized Ca^2+^ fluorescent signals (dF/F) according to the formula: dF/F = (F_i_-F_0_) / F_0_, where i represented each movie frame. The resulting movies were then used to extract fluorescent signals associated with individual cells using the Principal and Independent Component Analysis (PCA-ICA). The ROIs identified by PCA-ICA were further examined by a skilled experimenter based on dF/F traces and morphology. To be considered as a putative neuron, the Ca^2+^ waveform should exhibit a fast-rising onset followed by slow decaying signal. Additionally, the ROI morphology should have a spherical shape with high intensity, indicative of a cell body, rather than appearing as a small dot or line which could suggest non-specific, spine or axon signals (*67*). The dF/F of Ca^2+^ signal for each cell was further standardized as z-score using the formula: z-score = (dF/F-mean)/standard deviation.

*Ca^2+^ activity analyses:* To identify Ca^2+^ activity associated with foot-shocks in the CFC trainings, freezing behaviors during CFC memory tests or with immobile behaviors during the tail suspension test, we aligned single neuronal responses to the start and end of these behaviors. Ca^2+^ activity within a time-window of -5 sec to 5 sec around each alignment was averaged. The mean responses of all neurons were then clustered into groups using k-means clustering as previously described (*68*). In the CFC training phases, we defined neurons whose activity increased upon the onset of the foot-shocks as activated neurons and decreased upon the onset of the foot-shocks as inhibited neurons. The others were defined as neutral neurons. In the memory tests and tail suspension test, neurons that showed an increase in activity at the start of behavior and a decrease at the end were classified as activated neurons, while those that showed a decrease at the start and an increase at the end were classified as inhibited neurons. The remaining neurons classified as neutral. No statistical z-score threshold was applied to determine activated or inhibited neurons. To quantify changes in Ca^2+^ activity, we compared the AUC per sec for the -5 sec to 0 sec window with that for the 0-to 5-sec window at the onset or offset of the behaviors as specified in the figures.

To quantify the overall Ca^2+^ activity during memory tests and tail suspension, we first re-scale the raw z score to 0 to 2, proportionally. We then calculate the relative frequency distribution of the averaged area under the curve (AUC) per sec of each cell during the first 180 sec of all the tests, and compared the averaged AUC per sec for each animal between tests.

To determine the proportion of “generalization cells” or “strength cells” that overlapped with the “depression cells” during tail suspension, we used a permutation test (shuffled times = 10, 000) to assess whether the observed number of overlapping cells were significantly different from what would be expected by chance (*68*).

**Retro-TRAP and RNA sequencing**

The procedure was adapted from previous reports (*36, 37*). AAV expressing an anti-GFP nanobody fused to RPL10a in a Cre-dependent manner (AAV8-EF1a-FLEX-NBL10, 1.01×10^12^ vg/ml, 100nl) was injected into the mPFC, and retrograde AAV expressing Cre recombinase fused to GFP (AAVretro-hSyn-Cre-GFP, 1.63×10^13^ vg/ml, 200nl) was injected into the BNST. Four weeks after surgery, the mice were randomly divided into three groups and received either a single CFC training, two CFC trainings in the same context or two CFC trainings in altered contexts. The mice were sacrificed 1 h after the last CFC training. Their brains were sliced into 1 mm sections on a brain matrix (RWD Life Science, Shenzhen, China) in ice-cold dissection buffer (2.6 mM KCl, 1.23 mM NaH_2_PO_4_, 26.2 mM NaHCO_3_, 5 mM kynurenic acid, 212.7 mM sucrose, 10 mM dextrose, 0.5 mM CaCl_2_, 1 mM MgCl_2_). The mPFC was dissected out using a 15G puncher. The mPFC from 6-7 mice were pooled together as one sample, and homogenized in 500 μL ice-cold homogenization buffer [10 mM HEPES (pH 7.4), 150 mM KCl, 5 mM MgCl_2_, 1% NP40, 1mM DTT(Sigma), 80 U/ml RNAase inhibitor (Thermo), halt protein & phosphatase inhibitor (Thermo), 125 mg/ml cycloheximide and 200 ng/ml rNB (ChromoTek)] in a glass homogenizer (Glas-Col, USA). The homogenate was centrifuged at 21,130 x g for 10 min at 4℃. The supernatant was collected. 20 μL of the supernatant was saved as the input sample, whereas the remaining was incubated with 5 μg GFP antibody (Thermo Fisher, G10362) for 4 h at 4℃ with gentle mixing. Afterwards, 100 μL protein A conjugated Dynabeads (Thermo) was added into each sample and incubated for another hour at 4℃ with gentle mixing. After washes to remove nonspecific binding, the beads were incubated with 350 μL RLT buffer supplemented with β-mercaptoethanol (Qiagen) to dissociate immunoprecipitated (IP) RNA from the protein complex. IP and Input RNA was isolated using the RNAeasy Micro Kit following the manufacturer’s protocol (Qiagen, Hilden, Germany). The RNA was amplified and cDNA library was constructed using SMARTer Stranded Total RNA-Seq Kit v2 - Pico Input Mammalian (Takara Bio, USA) following manufacturer’s protocol. The cDNA samples were submitted to the Novogene (Beijing, China) for quality examination, and sequenced by an Illumina NovaSeq 6000 system as paired-end 150 bp reads.

Raw data from the sequencing underwent quality control. The data were pre-processed to acquire clean reads, mapped to the mouse reference genome and quantified using the Hisat2 (version 2.0.5) and featureCounts (version 1.5.0-p3). To determine the efficiency of the retro-TRAP, we analyzed relative expression levels of neuronal marker genes and glia marker genes comparing IP and Input samples using DESeq2 package (version 1.40.1) in R. Then focusing on the IP samples, Principal component analysis (PCA) was conducted using DESeq2 (version 1.40.1) and ggplot2 (version 3.4.2) packages in R/Bioconductor, and limma package (3.56.1) was used to remove experimental batch effect. Pairwise differential expression genes (DEGs) analysis was further carried out using DESeq2 (version 1.40.1). The transcripts were considered as DEGs with a cutoff as *p* < 0.01. Venn plot, heatmaps and volcano plots were generated using VennDiagram (version 1.7.3), pheatmap (version 1.0.12) and graphics (version 4.3.0) packages in R. Function enrichment of DEGs was analyzed using clusterProfiler (version 4.8.1) package in R/Bioconductor. The RNA sequencing data generated for this study have been deposited to the GEO repository under accession number GSE248449.

**Statistics and reproducibility**

Data analyses were performed with Prism v.8 software (GraphPad Software, San Diego, California). The Shapiro-Wilk test and the Kolmogorov-Smirnov test were conducted to evaluate the normality of the data distribution. Outliers were identified by the ROUT test with a Q value cutoff as 5%, and excluded from statistical analysis. Sample size was determined based on previous studies or pilot experiments, but not formally tested. All the experiments were repeated independently in at least three cohorts of animals to ensure the replicability of the findings and most experiments were conducted by at least two experimenters. The data were pooled to achieve the reported final sample size. For experiments with two groups, the two-tailed unpaired Student’s *t* test was used to compare the difference for normally-distributed datasets, and the Mann-Whitney test was used for datasets that were non-normally distributed. The one-way analysis of variance (ANOVA) or two-way ANOVA followed by *post hoc* tests as specified in the related figure legends were performed for multiple comparisons. The significance level was set to *p* < 0.05.

**Data availability**

The RNA sequencing data generated for this study have been deposited to the GEO repository under accession number GSE248449. All other data are included in the main text and supplementary materials of this article. Further information of this study is available upon reasonable request from the corresponding authors.

**Supplementary Figure Legends**

**Supplementary Figure 1. Correlations between memory performances and depression-like behaviors.** G index: the generalization index calculated as % freezing during G test divided by % freezing during F test. F test: the percentage of time spent in freezing during the memory strength test. G test: the percentage of time spent in freezing during the memory generalization test. NSF: the latency to eat in novelty-suppressed feeding test. ST: the grooming duration in splash test. TS: the immobile time during tail suspension test. SIT-DI: the differential index (DI) in social interaction test. DI was calculated as (investigation time of a social mouse – investigation time of toy mouse) / (total investigation time of the social mouse and the toy mouse). n = 24. Color and size of the circles indicate the nominal *p* value and the absolute *r* value from Pearson’s correlation analyses. * *p* < 0.05, ** *p* < 0.01, *** *p* < 0.001.

**Supplementary Figure 2. Changes in c-FOS expression in various upstream regions of the BNST by fear memory strength (F test) *versus* generalization (G test).** (**A**) Schematic of the hypothesis that BNST integrates different inputs to modulate freezing during memory tests, and cholera toxin B (CTB) injection into BNST and co-staining with c-FOS to mark activated input neurons. A representative image of CTB infusion into BNST. Scale bar, 350 µm. (**B, D**) Schematics of the experiments. (**C, E**) Fold changes in c-FOS expression in different upstream regions of the BNST by the memory generalization test (G test, **C,** n = 8-11 per group) and the memory strength test (F test, **E,** n = 6-8 per group), normalized to the mean value for the “1xTr” group in each region. (**F**) Representative images of c-FOS staining in the CTB-labelled BNST-projecting neurons in various upstream brain regions by the F test. Scale bar, 40 µm. (**G**) Fold changes in c-FOS expression in the CTB^+^ BNST-projecting neurons in upstream regions by the F test, normalized to the mean value for the “1xTr” group in each region. n = 5-7 per group. Data are presented as mean ± s.e.m. and analyzed by multiple *t* tests. * *p* < 0.05, *** *p* < 0.001, **** *p* < 0.0001.

**Supplementary Figure 3. Coronal diagrams summarizing implant placements.**

**Supplementary Figure 4. Modulation of the mPFC^BNST^ neuronal calcium activity by contextual fear conditioning (CFC).** (**A**) Quantification of the number of neurons that showed significant changes in activity in response to foot-shocks during the first (CFC1) and the second (CFC2) trainings in the same (2xTr same) or different contexts (2xTr altered). (**B**) Heatmaps of the calcium activity of the shock-activated and inhibited cells during CFC1 and CFC2 training sessions. (**C-F**) Averaged curves and quantification of the difference in area under the curve per second (AUC/sec) between 20 sec pre- and post- foot-shocks for the shock-activated (**C, E**) and inhibited (**D, F**) cells during CFC1 and CFC2 training sessions in the “2xTr same” (**C-D**) and “2xTr altered” (**E-F**) groups. “2xTr same” group: n = 6 animals, 383 (CFC 1) and 360 (CFC 2) shock-activated cells; 256 (CFC 1) and 276 (CFC 2) shock-inhibited cells. “2xTr altered” group: n = 9 animals, 473 (CFC 1) and 441 (CFC 2) shock-activated cells; 280 (CFC 1) and 273 (CFC 2) shock-inhibited cells. Data are presented as box-and-whiskers graphs, with the box extends from the 25^th^ to 75^th^ percentile, the line in the middle of the box is plotted at the median and the whiskers extend from 10 to 90 percentiles. Data are analyzed by two-way ANOVA followed by Sidak’s p*ost hoc* test. * *p* < 0.05, ** *p* < 0.01, *** *p* < 0.001, **** *p* < 0.0001.

**Supplementary Figure 5. Quantification of the differences in area under the curve per second (AUC/sec) of mPFC^BNST^ neuronal calcium activity, comparing 5 sec before and after the behavioral events.** (**A**) Quantification of AUC/sec during behavioral tests after 2xTr same training. (**B**) Quantification of AUC/sec during behavioral tests after 2xTr altered training. F test: memory strength test in the same context as used during training. G test: memory generalization test in the modified context. TS: tail suspension test for assessing depression-like behaviors. ** *p* < 0.01, *** *p* < 0.001, **** *p* < 0.0001.

**Supplementary Figure 6. Connectivity of the mPFC-to-BNST projections.** (**A-B**) Schematic of cholera toxin B (CTB) infusion into the BNST, and representative images of CTB in the BNST and in the mPFC. Scale bar, 350 µm. (**C**) Quantification of CTB-labelled neurons in different subregions as a percentage of total mPFC^BNST^ neurons. (**D**) Representative images of CTB-labelled mPFC^BNST^ neurons (red) in the mPFC, co-stained with CaMKII or GABA (grey). Scale bar, 100 µm. (**E**) Quantification of the percentage of CaMKII^+^ and GABA^+^ neurons in the mPFC^BNST^ neurons. (**F**) Schematics of AAV infusions for labeling inputs to the mPFC^BNST^ neurons, and representative images of mPFC^BNST^ neurons identified as starter cells co-expressing EGFP and dsRed (yellow). Scale bar, 50 μm. (**G**) Representative images of dsRed-labelled input neurons targeting the mPFC^BNST^ neurons. Scale bar, 300 μm. (**H**) Quantification of the amount of input cells in each region as a percentage of total input cells. n = 5 per region. (**I**) Schematics of AAV infusions and a representative image of AAV-mediated mCherry expression in the mPFC. Scale bar, 200 µm. (**J**) Representative images of mCherry-labelled mPFC projections in the BNST. Scale bar, 350 µm. (**K**) Quantification of the area covered by mPFC projections in different BNST subregions as a percentage of the overall BNST. (**L**) Schematics of AAV infusions for labeling BNST^mPFC^ neurons. (**M**) Representative images of the BNST^mPFC^ neurons (red) co-stained with CaMKII or GABA (grey). Scale bar, 100 μm. (**N**) Quantification of the percentage of CaMKII^+^ and GABA^+^ neurons in all mCherry-labelled BNST^mPFC^ neurons. (**O**) Representative images of mCherry-labelled BNST^mPFC^ projections in various downstream regions. Scale bar, 300 μm. (**P**) Quantification of BNST fiber-covered area as a percentage of the area of each downstream region. n = 4-5 per region. Data are presented as mean ± s.e.m.

**Supplementary Figure 7. Activation of the mPFC-to-BNST projections promotes depression-like behaviors at 2 weeks after training, related to Figure 5.** (**A, F**) Schematics of AAV infusion and experimental timelines. (**B, G**) Quantification of the latency to eat in novelty-suppressed feeding (NSF) test. (**C, H**) Quantification of the grooming duration in splash test (ST). (**D, I**) Quantification of the immobile time during tail suspension (TS) test. (**E, J**) Quantification of the exploration time and differential index (DI) in social interaction test (SIT). **B-E**, changes in behaviors by chemogenetic inhibition of the mPFC-to-BNST projections after “2xTr altered”. n = 8-11 per group. **G-J**, changes in behaviors by optogenetic activation of the mPFC-to-BNST projections after “2xTr same”. n = 6-10 per group. Data are presented as mean ± s.e.m. and analyzed by *t* tests and two-way ANOVA followed by Sidak’s *post hoc* test. * *p* < 0.05, ** *p* < 0.01, *** *p* < 0.001, **** *p* < 0.0001.

**Supplementary Table 1. List of brain regions analyzed in c-FOS network analysis.**

| **Region** | **Abbreviation** | **#** |
| --- | --- | --- |
| medial prefrontal cortex | mPFC | 1 |
| anterior cingulate cortex | ACC | 2 |
| lateral orbital cortex | LO | 3 |
| nucleus accumbens, core | NAcC | 4 |
| nucleus accumbens, shell | NAcS | 5 |
| agranular insular cortex | AI | 6 |
| bed nucleus of stria terminalis | BNST | 7 |
| reuniens thalamic nucleus | Re | 8 |
| paraventricular hypothalamic nucleus | PVN | 9 |
| lateral hypothalamus | LH | 10 |
| dorsomedial hypothalamus | DMH | 11 |
| ventromedial hypothalamus | VMH | 12 |
| arcuate hypothalamic nucleus | ARC | 13 |
| paraventricular thalamus | PVT | 14 |
| lateral habenula | LHb | 15 |
| basolateral amygdala | BLA | 16 |
| central amygdala | CeA | 17 |
| cortical amygdala | CoA | 18 |
| field CA1 of dorsal hippocampus | dCA1 | 19 |
| field CA3 of dorsal hippocampus | dCA3 | 20 |
| dentate gyrus, dorsal part | dDG | 21 |
| field CA1 of ventral hippocampus | vCA1 | 22 |
| ventral subiculum | vSUB | 23 |
| lateral entorhinal cortex | LEnt | 24 |
| perirhinal cortex | PRh | 25 |
| entorhinal cortex | Ect | 26 |
| retrosplenial granular cortex | RSG | 27 |
| ventral tegmental area | VTA | 28 |
| lateral periaqueductal gray | lPAG | 29 |
| medial periaqueductal gray | mPAG | 30 |
