## Supplementary figures and images for "Generalization but not strengthening of negative memories drives the development of depression-like behaviors"

### Supplemental figure1

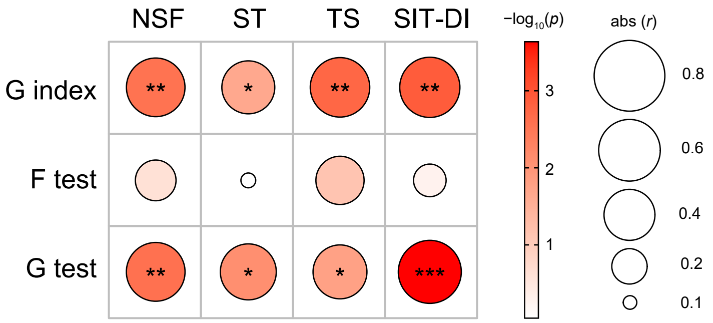

### Supplemental figure2

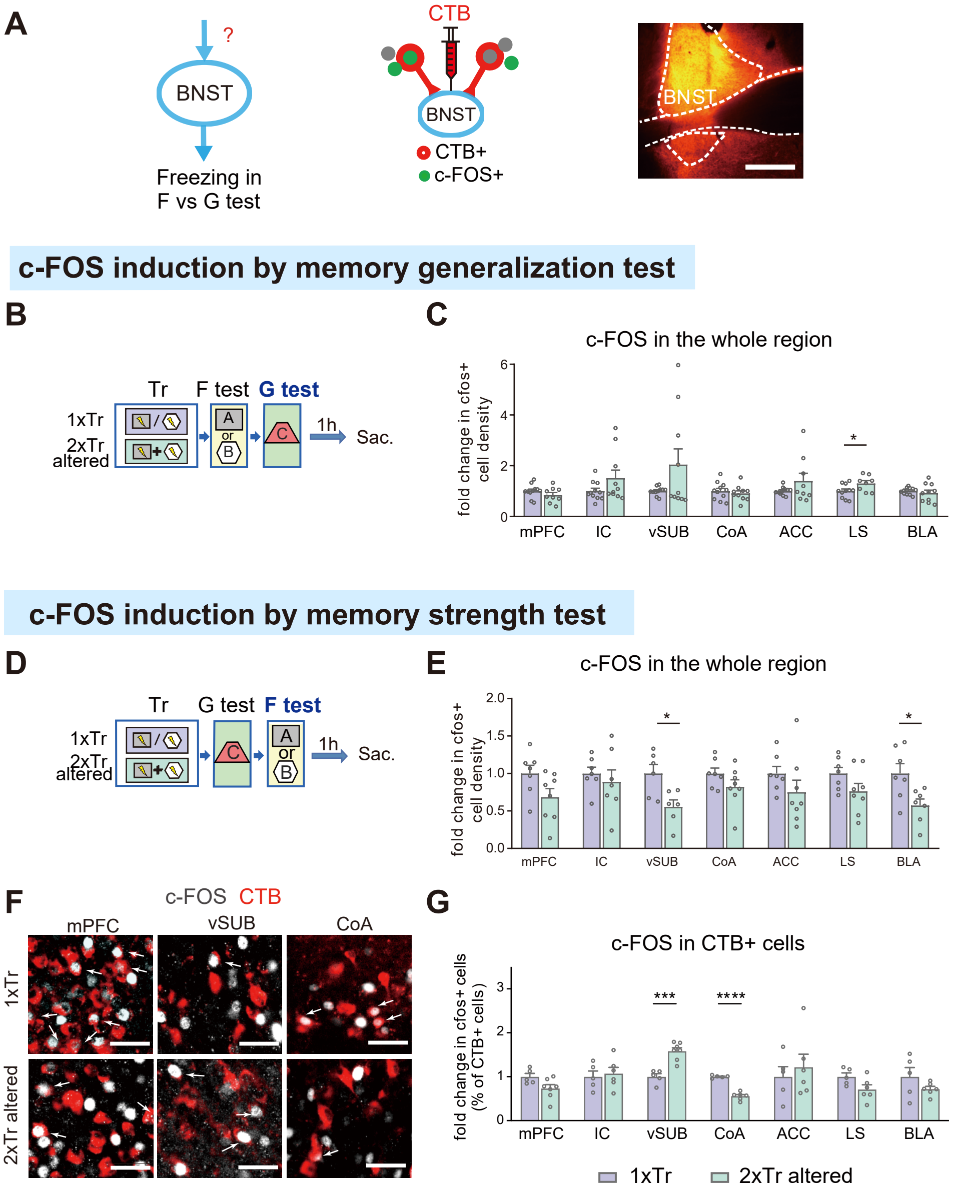

### Supplemental figure3

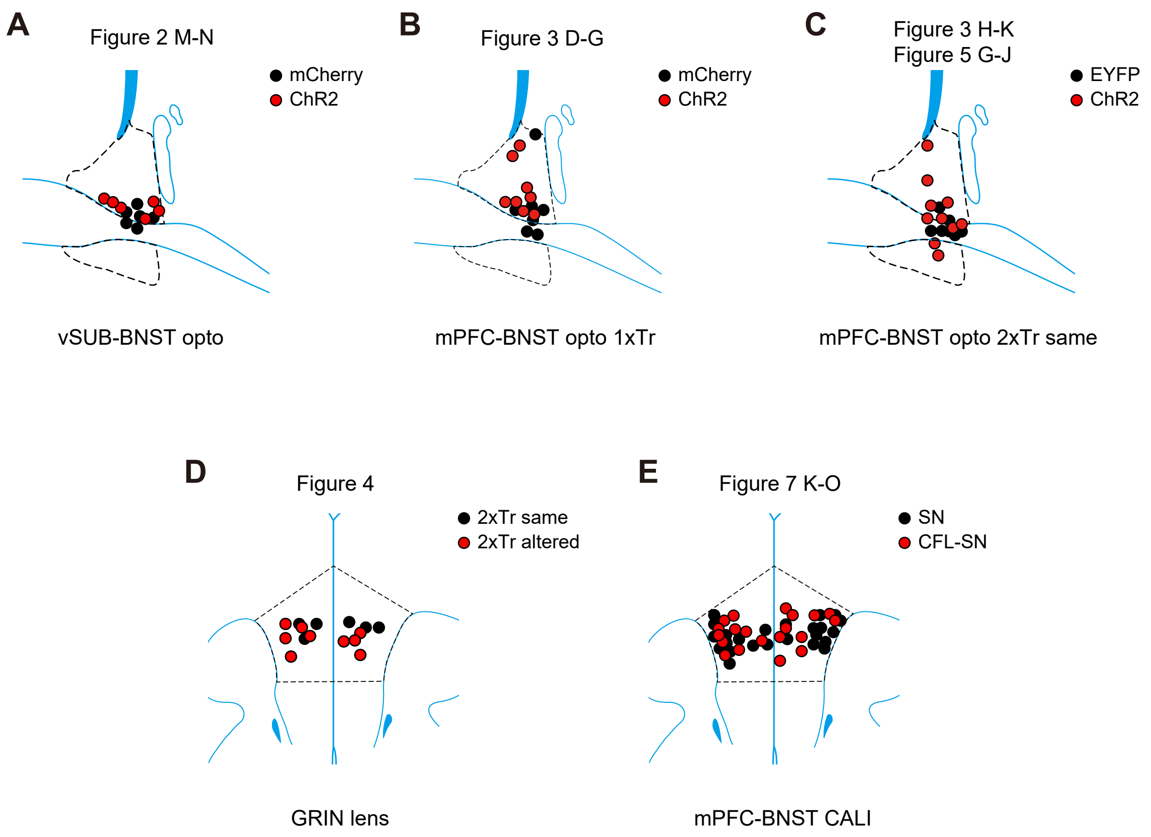

### Supplemental figure4

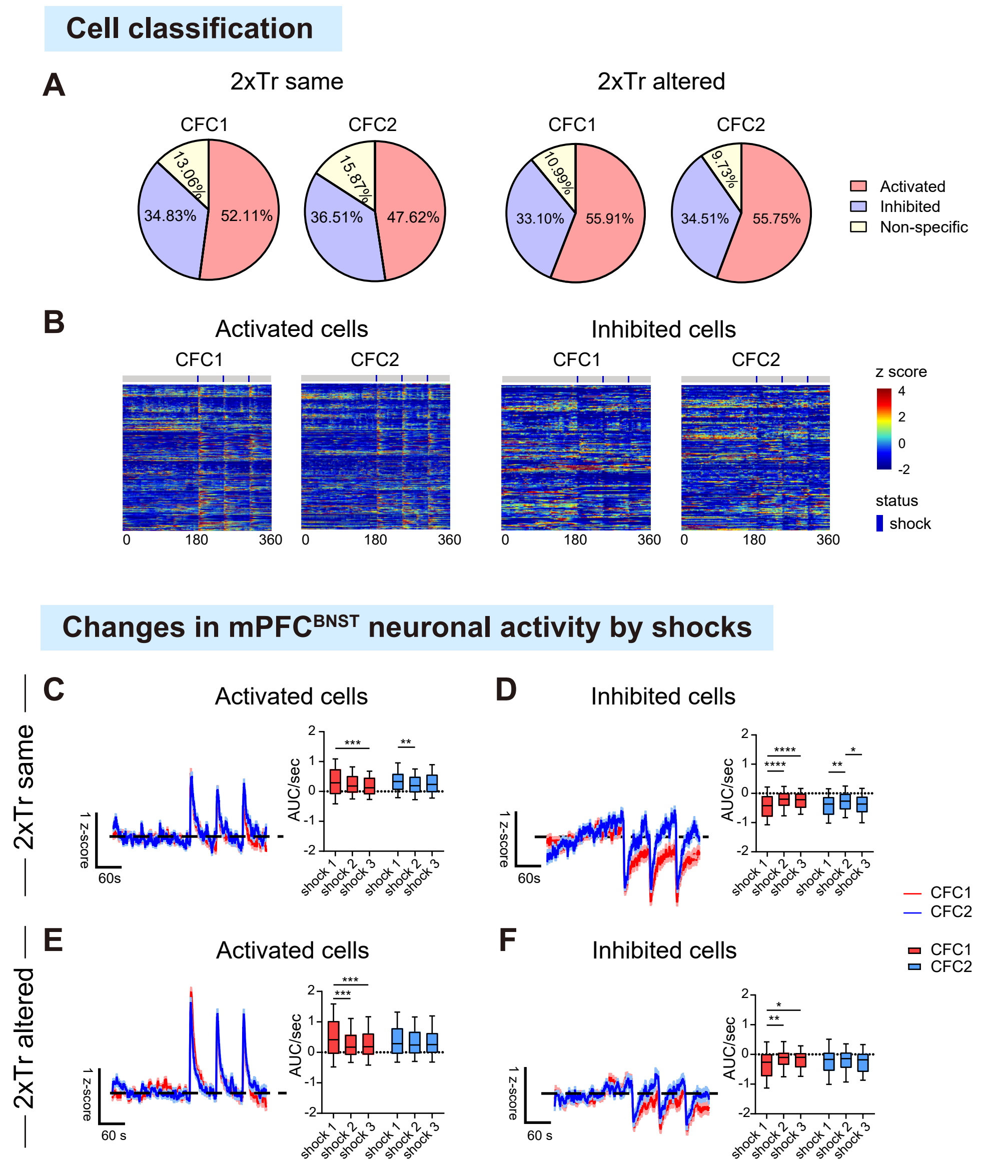

### Supplemental figure5

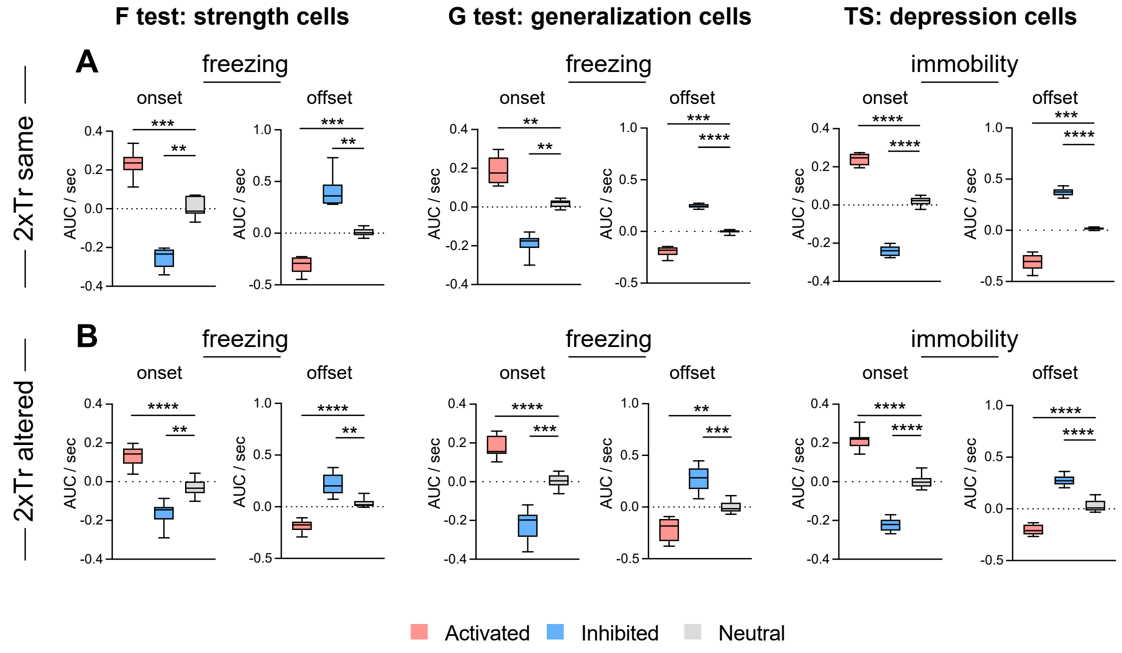

### Supplemental figure6

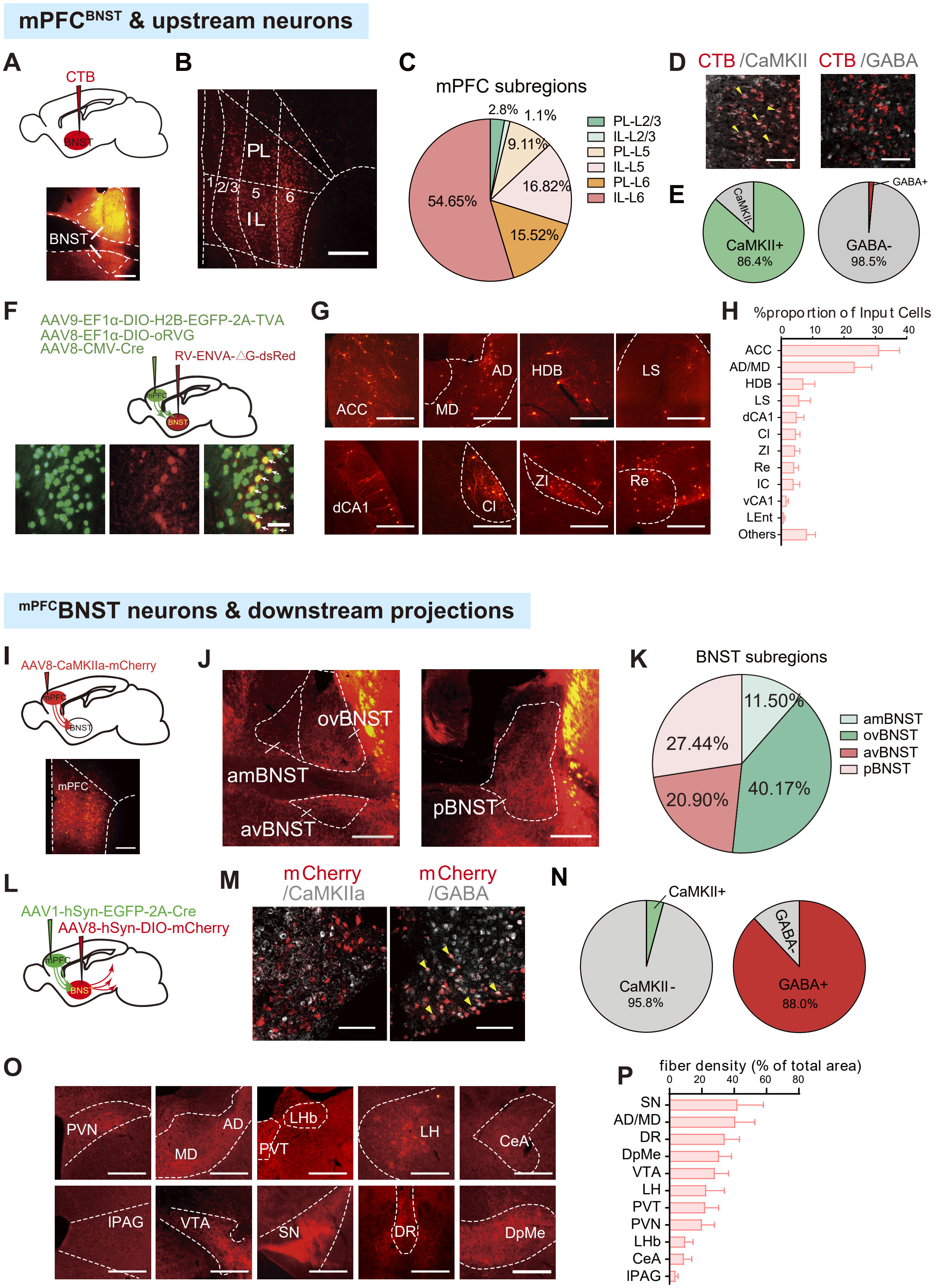

### Supplemental figure7

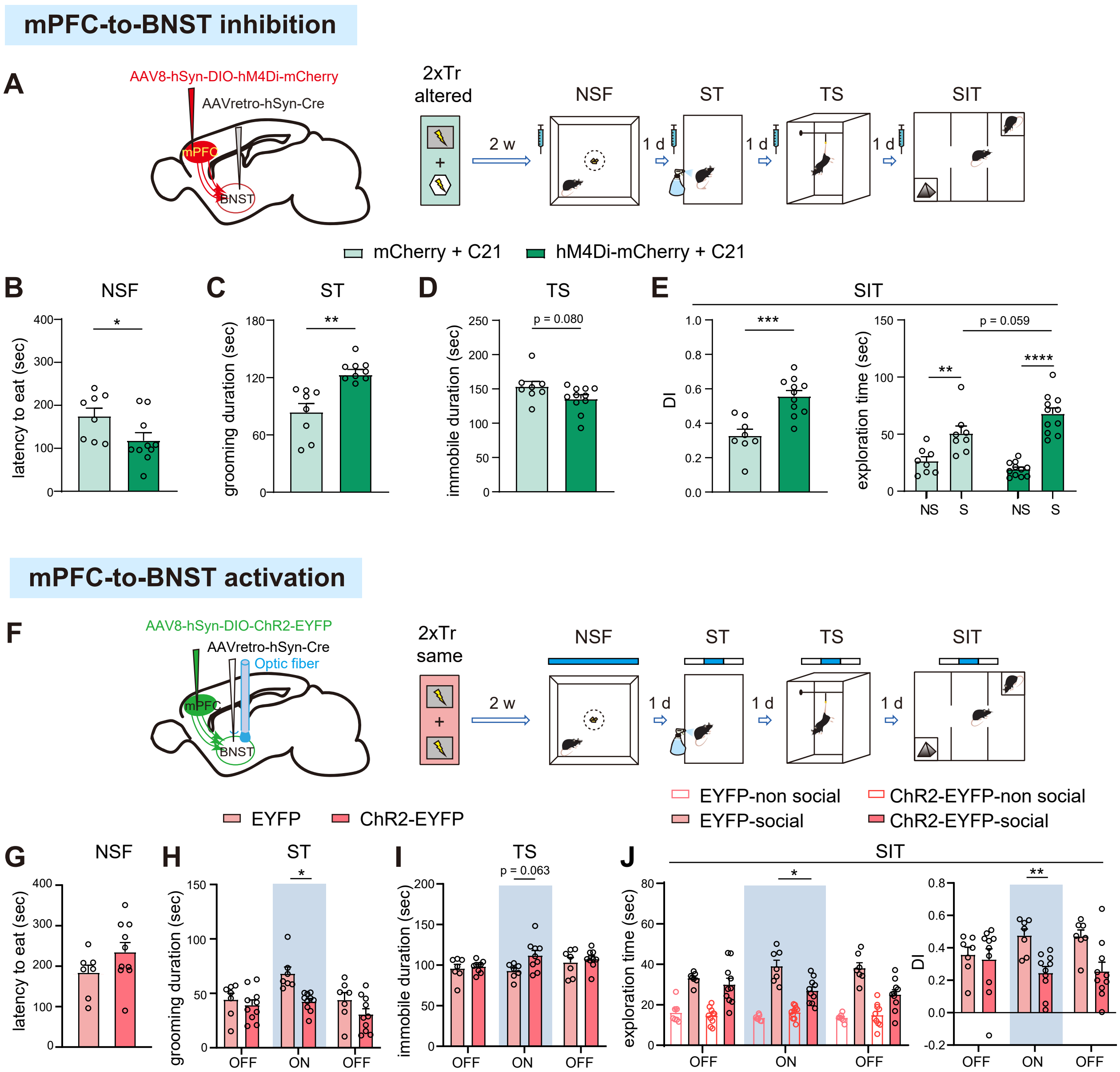
